# Macropinocytosis controls metabolic stress-driven CAF subtype identity in pancreatic cancer

**DOI:** 10.1101/2024.11.29.625709

**Authors:** Yijuan Zhang, Li Ling, Swetha Maganti, Jennifer L. Hope, Cheska Marie Galapate, Florent Carrette, Karen Duong-Polk, Anindya Bagchi, David A. Scott, Andrew M. Lowy, Linda M. Bradley, Cosimo Commisso

## Abstract

Pancreatic ductal adenocarcinoma (PDAC) tumors are deficient in glutamine, an amino acid that tumor cells and CAFs use to sustain their fitness. In PDAC, both cell types stimulate macropinocytosis as an adaptive response to glutamine depletion. CAFs play a critical role in sculpting the tumor microenvironment, yet how adaptations to metabolic stress impact the stromal architecture remains elusive. In this study, we find that macropinocytosis functions to control CAF subtype identity when glutamine is limiting. Our data demonstrate that metabolic stress leads to an intrinsic inflammatory CAF (iCAF) program driven by MEK/ERK signaling. Utilizing *in vivo* models, we find that blocking macropinocytosis alters CAF subtypes and reorganizes the tumor stroma. Importantly, these changes in stromal architecture can be exploited to sensitize PDAC to immunotherapy and chemotherapy. Our findings demonstrate that metabolic stress plays a role in shaping the tumor microenvironment, and that this attribute can be harnessed for therapeutic impact.

**Graphical Abstract:** 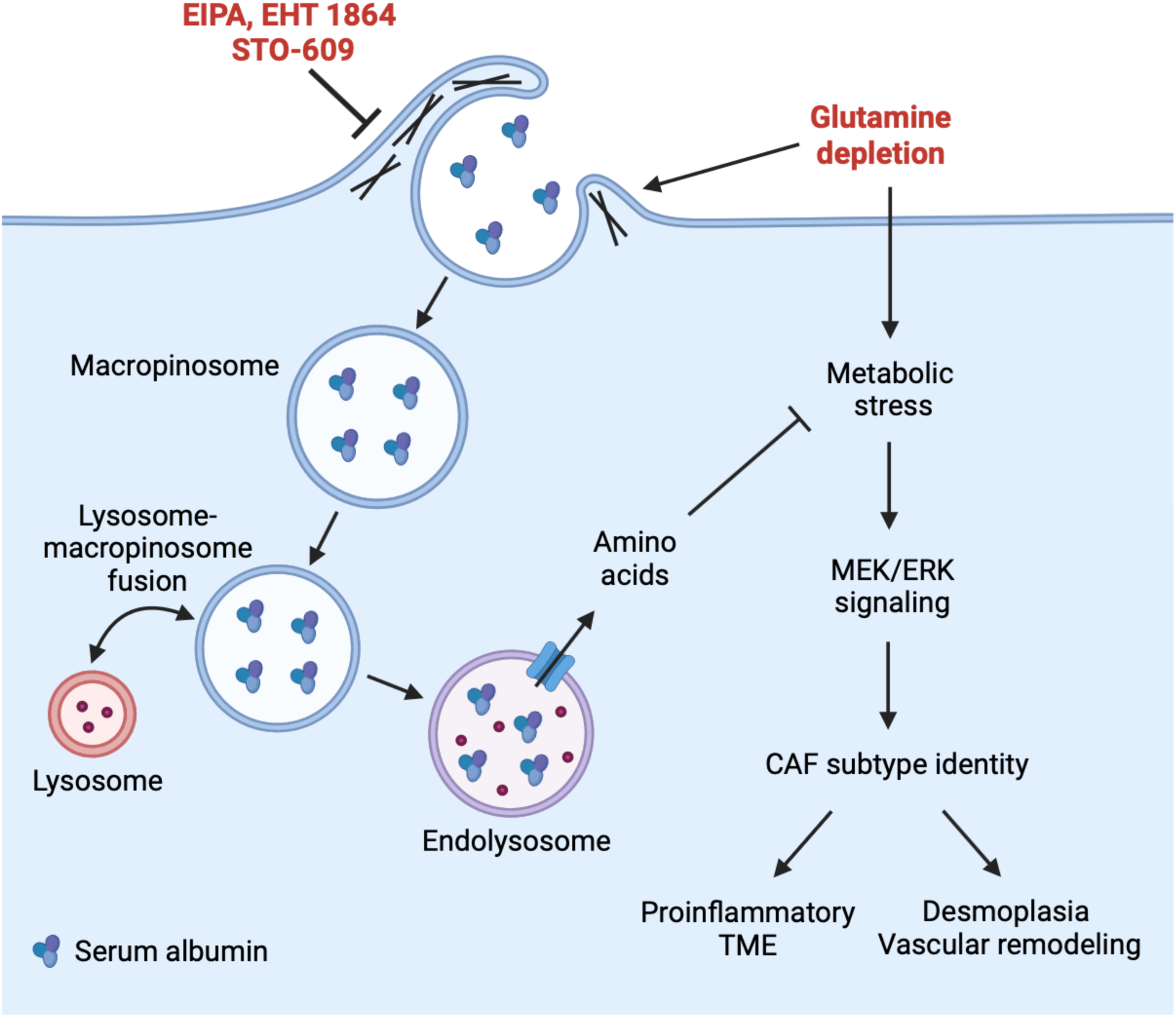

## Introduction

Cancer-associated fibroblasts (CAFs) are a prominent feature of the tumor microenvironment (TME) in pancreatic ductal adenocarcinoma (PDAC), and they have multiple roles in modulating the activity of both cancer cells and tumor-infiltrating immune cells^1–5^. CAFs have been demonstrated to support PDAC tumor growth through the secretion of cytokines and growth factors, providing metabolites, and promoting immune evasion^6–9^. The extent of heterogeneity displayed by pancreatic CAFs is becoming greatly appreciated, and the functionality of the different CAF subtypes is thought to vary, with examples of both tumor-promoting and tumor-suppressive roles in different contexts^3,10–18^. The majority of CAFs in PDAC tumors are classified as myofibroblasts (myCAFs), which strongly express alpha smooth muscle actin (α-SMA) and contribute to the desmoplastic TME through the deposition of collagen and other extracellular matrix (ECM) proteins^19,20^. This dense stromal reaction is thought to serve as a barrier that impedes both immune cell infiltration and the delivery of chemotherapeutic drugs. Another major subtype of CAFs found within the pancreatic TME are the inflammatory CAFs (iCAFs), which are a subset of IL-1-induced CAFs that express proinflammatory cytokines and chemokines, including IL-6, CXCL-1 and CXCL-12^14^. Validation by single-cell RNA sequencing has established specific markers for myCAFs, iCAFs and other CAF subtypes found within PDAC tumors^15^. Recent studies have revealed that cellular plasticity is an important feature of CAFs. For example, myCAFs and iCAFs can be interconverted under different extrinsic stimuli, such as Hedgehog, TGF-β or IL-1/TNF-α signaling^3,20,21^. Also, the extrinsic IL-1-induced iCAF phenotype can be potentiated by hypoxia signaling or the activation of p38 MAPK in cancer cells^22–24^. Intratumoral nutrient deprivation, such as amino acid deficiency, has been demonstrated to induce metabolic stress in PDAC tumors^25–27^. What is unclear is whether this metabolic stress caused by amino acid scarcity can modulate CAF plasticity, heterogeneity, or identity. Moreover, if metabolic stress does impact CAF subtypes, it would be beneficial to determine whether such effects can be harnessed to sensitize PDAC to immunotherapy or chemotherapy.

A biological property of CAFs found in the pancreatic TME is the ability to carry out macropinocytosis, an actin-dependent endocytic pathway that mediates the non-selective uptake of extracellular fluid^7^. In the context of PDAC, macropinocytosis was first described as an important feature of the tumor cells that drives the uptake of extracellular serum albumin, which, when targeted for lysosome-mediated degradation, serves as a rich source of protein-derived amino acids that the tumor cells use to sustain their proliferative capacity despite the nutrient-depleted state of the pancreatic TME^27–29^. Interestingly, recent evidence points to the notion that macropinocytosis is an integral aspect of the metabolic rewiring that occurs throughout PDAC tumors, as it is employed by both the cancer cells and the CAFs to support their cellular fitness^7,26^. An additional tumor-promoting feature of macropinocytic stimulation in the CAFs is that the excess protein-derived amino acids that are produced by this catabolic pathway can be secreted, nourishing the tumor cells. What remains unclear is whether macropinocytic induction in the CAFs serves to modulate responses to metabolic stress to support and maintain CAF heterogeneity within the TME. Moreover, it is unclear how interfering with macropinocytosis in both the tumor cells and the CAFs might alter PDAC progression and the stromal landscape.

In this study, we find that metabolic stress can trigger an intrinsic inflammatory CAF program in PDAC. Using multiple inhibitors of macropinocytosis, we show that enhanced metabolic stress in the context of glutamine depletion leads to an increase in iCAF markers and a concomitant reduction of myCAF markers, indicating that macropinocytosis functions in maintaining CAF subtype identity. Dissection of the underlying molecular mechanism reveals that MEK/ERK signaling is the key regulator driving this process. We find that the *in vivo* inhibition of macropinocytosis with 5-(N-ethyl-N-isopropyl)-amiloride (EIPA), a specific macropinocytosis inhibitor, causes the suppression of tumor growth that is accompanied by changes in the stromal architecture. We observe that EIPA leads to an enrichment of iCAFs and a diminishment of myCAFs within the PDAC tumors, which is associated with the specific enhancement of tumor-infiltrating CD4^+^ and CD8^+^ T cells and a proinflammatory TME. Importantly, combining macropinocytosis inhibition with immune checkpoint blockade suppresses PDAC progression. Lastly, we establish that the decrease in collagen deposition and expanded vasculature associated with reductions in the myCAF population allow for the improved delivery of chemotherapy to the tumors, resulting in macropinocytosis inhibition synergistically sensitizing PDAC tumors to gemcitabine. Overall, our work identifies an intrinsic regulatory program that controls CAF heterogeneity in response to metabolic stress and points to the potential application of macropinocytosis inhibitors as a strategy to sensitize PDAC to immunotherapy or chemotherapy.

## Results

### Metabolic stress induces inflammatory markers in pancreatic CAFs

We and others have shown that glutamine levels are depleted in PDAC in the context of human clinical specimens, xenograft tumors and murine syngeneic orthotopic tumors^25–27^. These previous analyses were performed using bulk tumor metabolomics; therefore, we sought to determine whether tumor interstitial fluid (TIF) was also depleted of glutamine. To do this, we isolated TIF using established procedures^30^ from both orthotopic and subcutaneous syngeneic tumors. We then quantified polar metabolites utilizing gas chromatography–mass spectrometry (GC-MS) and compared amino acid levels in the TIF relative to the plasma. In line with previous reports^30^, we found that the TIF was depleted of tryptophan, but enriched in aspartate, glutamate, glycine and ornithine (Figure S1A and S1B). We also found that glutamine was significantly depleted from the TIF relative to the plasma in both tumor models (Figure 1A). We previously reported that varying glutamine levels within the tumors has the ability to control macropinocytosis in both PDAC cells and CAFs, allowing both cell types to dial-up or dial-down this uptake pathway as needed to acquire protein-derived amino acids and maintain their cellular fitness^7,26–29^. CAF heterogeneity and plasticity has recently become a widely appreciated aspect of tumor progression, especially in PDAC ^3,11,12,14,15,20^. It is not clear how the metabolic stress caused by glutamine depletion within the tumors might influence CAF identity. Moreover, whether amino acids provided by the macropinocytosis pathway in response to glutamine stress play a role in maintaining or regulating CAF subtypes has never been explored. To start to answer these questions, we isolated CAFs from murine PDAC tumors that were positive for fibroblast specific protein 1 (FSP1) and podoplanin (PDPN), both established CAF markers (Figure S1C). We also verified that the isolated cells were negative for E-Cadherin, an epithelial cell marker, and positive for α-SMA, a myCAF marker that all CAFs express in culture (Figure S1D). In these murine CAFs, we found that macropinocytosis dampens the metabolic stress caused by low levels of glutamine because treatment with EIPA, a specific inhibitor of macropinocytosis, further increased the expression of Sestrin 2 (SESN2), a marker of amino acid stress (Figures 1B-1E)^25,26,31^. These data indicated that glutamine deprivation combined with macropinocytosis inhibition can lead to enhanced amino acid stress relative to either individual condition on its own.

**Figure 1.**
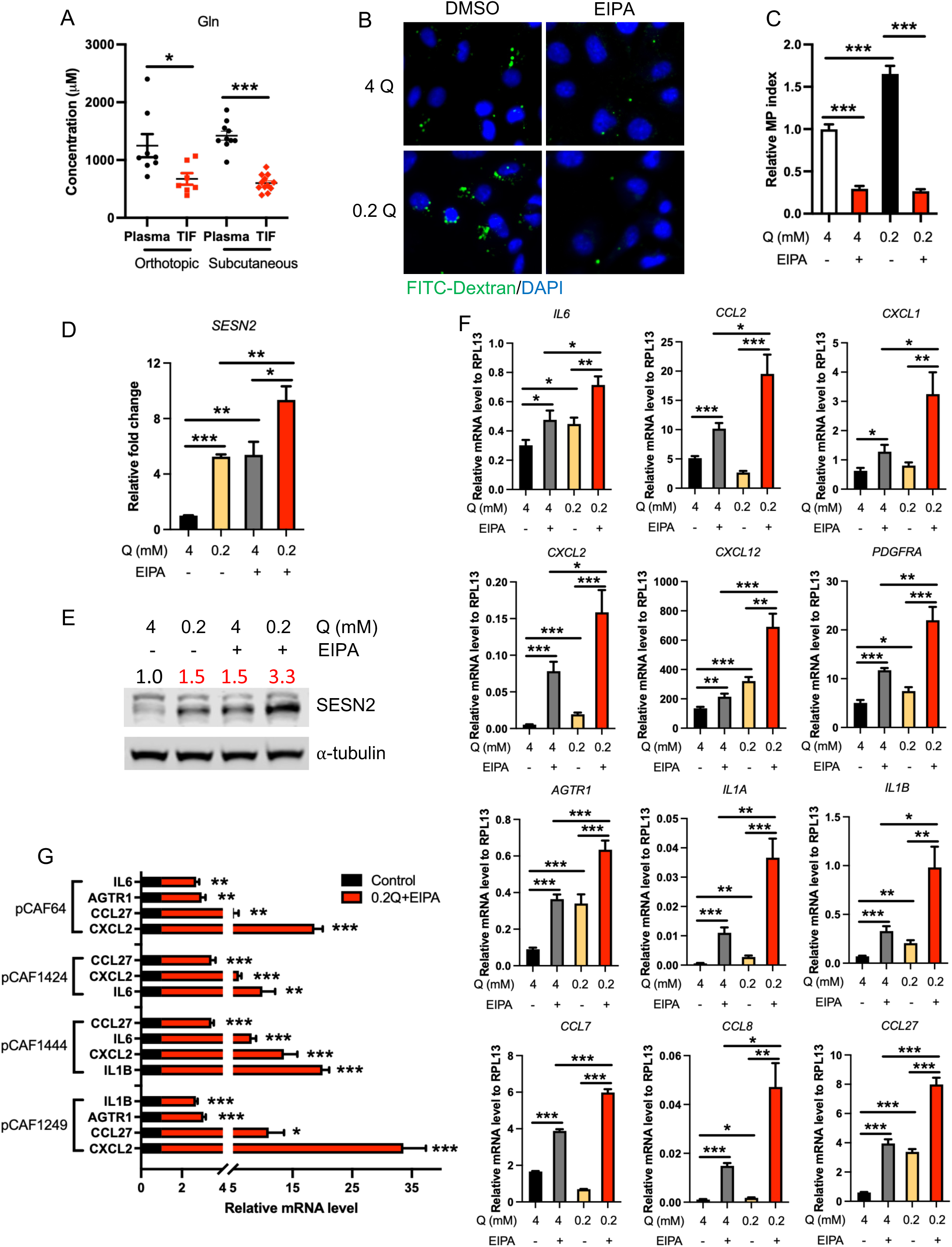
Glutamine deprivation and macropinocytosis cooperate to control metabolic stress and CAF subtype identity. (A) Quantitation of glutamine levels in plasma and tumor interstitial fluid (TIF) that were isolated from orthotopic or subcutaneous KPC tumors. Data are shown as mean ± SEM, unpaired t-test. *p<0.05, ***p<0.001. (B and C) Representative images (B) and quantification (C) of macropinocytic uptake of FITC-labeled dextran in CAFs treated with 4 mM or 0.2 mM glutamine (Q) with or without the presence of 25 μM EIPA for 24 hours. Representative data of three independent experiments are shown as mean ± SEM, unpaired t-test. ***p<0.001. Scale bar: 20 μm. (D and E) Mouse CAFs were treated with 4 mM or 0.2 mM glutamine (Q) with or without the presence of 25 μM EIPA for 2 or 4 hours. The mRNA (D) and protein (E) levels of sestrin2 (SESN2) were analyzed by qPCR and western blot, respectively. Data are shown as mean ± SEM, one-way ANOVA. *p<0.05, **p<0.01, ***p<0.001. (F) Mouse CAFs were treated with 4 mM or 0.2 mM Q with or without the presence of 25 μM EIPA for 32 hours. The mRNA levels of iCAF-related markers were analyzed by qPCR and normalized to RPL13. Data are shown as mean ± SEM, one-way ANOVA and unpaired t-test. *p<0.05, **p<0.01, ***p<0.001. (G) Four primary PDAC patient-derived CAFs (pCAF) were treated with 0.2 mM Q with 25 μM EIPA or vehicle control for 48 hours. The mRNA levels of iCAF-related markers were analyzed by qPCR. Data are shown as mean ± SEM, unpaired t-test. *p<0.05, **p<0.01, ***p<0.001.

Using the *in vitro* cell-based system described above, we next evaluated whether glutamine depletion, EIPA or the combination could regulate the expression of an established panel of iCAF or myCAF markers^15,20^. Interestingly, we found that low glutamine and macropinocytosis inhibition cooperated to significantly induce the expression of several iCAF and related inflammatory markers, including IL6, CCL2, CXCL1, CXCL2, CXCL12, PDGFRA, AGTR1, IL-1A, IL-1B, CCL7, CCL8, and CCL27 (Figure 1F). Concomitantly, we found that low glutamine combined with macropinocytosis inhibition also decreased the expression of myCAF markers including ACTA2, MYL9, TAGLN, and TPM2 (Figure S1E). Also, the most abundant fibrillar collagens, such as COL1A1, COL1A2, and COL3A1, which are pan-CAF markers but are more highly expressed in myCAFs relative to iCAFs^15^, were also reduced (Figure S1F). We also assessed the effects of metabolic stress on CAFs isolated from human PDAC patients. Using patient-derived CAFs originating from four different patients, we confirmed that low glutamine and macropinocytosis inhibition significantly, but heterogeneously, increased the expression of IL6, CXCL2, IL1B, AGTR1, and CCL27 (Figure 1G). We next examined whether glutamine depletion combined with other macropinocytosis inhibitors could also induce the inflammatory markers in CAFs. Previously, we demonstrated that STO-609, a CaMKK2 inhibitor, and Rac inhibitors, could suppress macropinocytosis in patient-derived pancreatic CAFs^7^. We found that treatment with STO-609 or EHT 1864, a Rac inhibitor, significantly inhibited macropinocytosis in the murine CAFs (Figures S2A, S2B, S2D and S2E). We next examined the effects on the expression of the inflammatory markers and found that STO-609 and EHT 1864, like EIPA, significantly induced the expression of the iCAF markers (Figures S2C and S2F). Altogether, these data demonstrate that metabolic stress caused by a combination of low glutamine and macropinocytosis inhibition can induce an intrinsic inflammatory program in pancreatic CAFs. Moreover, our data suggest that part of the normal function of macropinocytic induction within the stroma is to support and maintain CAF subtype identity and heterogeneity.

### The intrinsic metabolic stress-induced iCAF phenotype is linked to protein scavenging and is reversible

We previously reported that macropinocytosis functions in CAFs to support CAF viability through the uptake of serum albumin and the subsequent production of protein-derived amino acids^7^. We hypothesized that if we supplemented the growth medium with high levels of serum albumin (4%), then the inflammatory CAF phenotype caused by metabolic stress might be reversed. We initially tested the extent to which EIPA could inhibit macropinocytosis when the CAFs are cultured in serum albumin as a nutrient source. We found that, as expected, serum albumin supplementation dampens the extent of macropinocytosis occurring in the CAFs under conditions where glutamine is restricted (Figures 2A and 2B). It should be noted that under these conditions, EIPA only reduced macropinocytic capacity by approximately 40-50%, suggesting that the residual macropinocytosis could be sufficient to provide enough serum albumin-derived amino acids to suppress the inflammatory phenotype. Next, we cultured CAFs in low glutamine with and without serum albumin supplementation and treated the cells with EIPA or vehicle prior to analyzing the effects on the expression of the iCAF markers. We found that serum albumin diminished the EIPA-driven increases in expression of the inflammatory markers (Figure 2C), which indicates that the intrinsic induction of the iCAF phenotype is directly attributable to protein catabolism.

**Figure 2.**
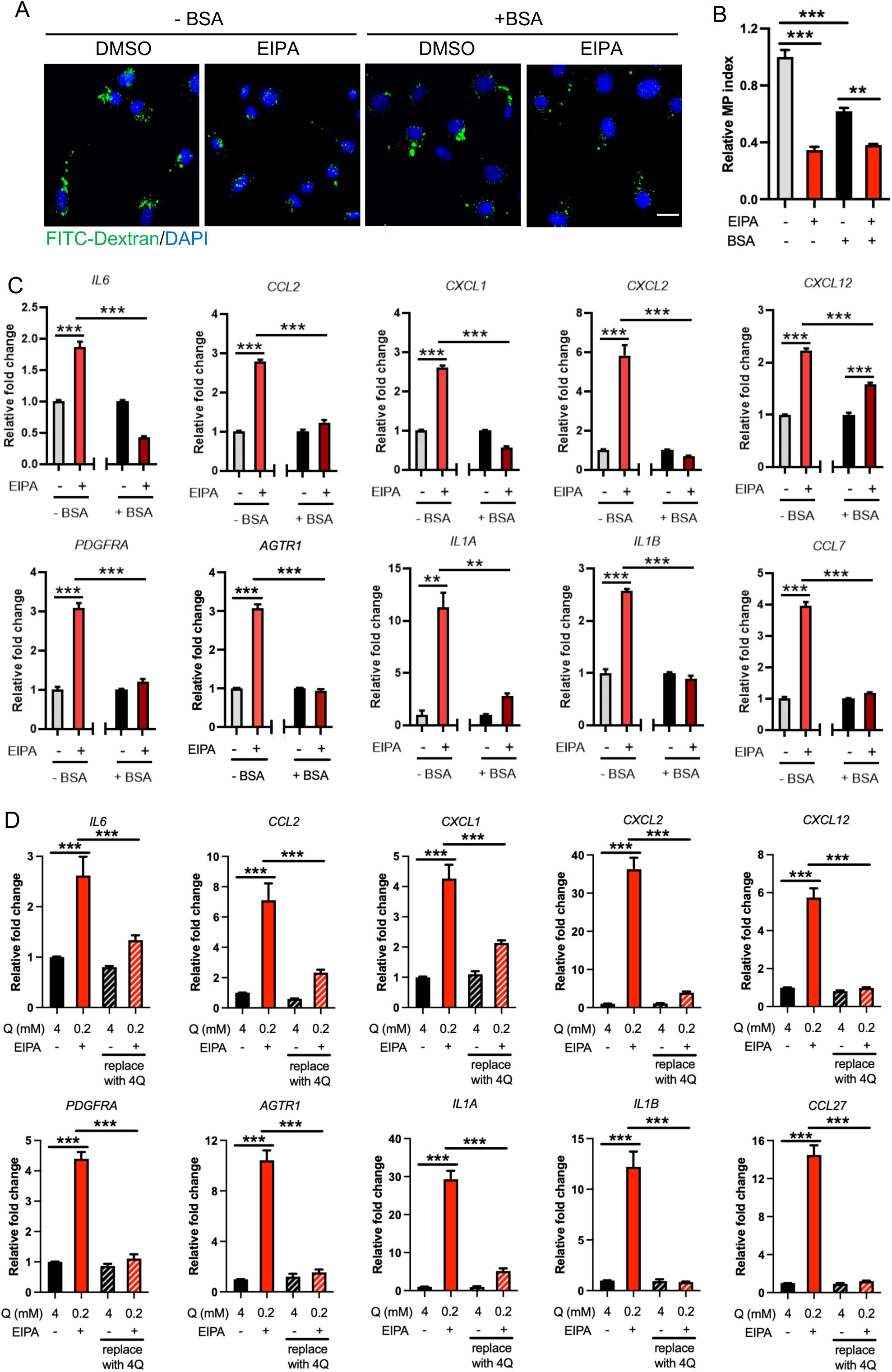
Induction of CAF inflammatory markers by metabolic stress is rescued by serum albumin supplementation and is reversible. (A and B) Representative images (A) and quantification (B) of macropinocytic uptake of FITC-labeled dextran in mouse CAFs treated with 0.2 mM Q with DMSO or EIPA with or without the presence of 40 mg/mL BSA. Representative data of three independent experiments are shown as mean ± SEM, One-way ANOVA. **p<0.01, ***p<0.001. Scale bar: 20 μm. (C) Mouse CAFs were treated with 0.2 mM Q with DMSO or EIPA with or without the presence of 4% BSA for 24 hours. The mRNA levels of iCAF-related markers were analyzed by qPCR. Data are shown as mean ± SEM, unpaired t-test. *p<0.05, **p<0.01, ***p<0.001. (D) Mouse CAFs were treated with 4 mM Q plus vehicle or 0.2 mM Q plus EIPA. 32 hours post treatment, one set of samples were collected as baseline and another set were washed with PBS and cultured with 4 mM Q medium for an additional 24 hours. Data are shown as mean ± SEM, unpaired t-test. ***p<0.001.

CAF subtype plasticity has been shown to be an important aspect of tumor progression in several cancer types^1,3,11,32^; therefore, we next sought to assess whether the induction of the inflammatory phenotype caused by metabolic stress was reversible. To do this, we cultured CAFs in low glutamine and EIPA for 32 hours, at which point the culture media was then replaced with glutamine replete media lacking EIPA for an additional 24 hours. We found that the expression of the iCAF markers was reversed after switching the cells back to regular media (Figure 2D). Also, we determined that the cell morphological changes associated with metabolic stress in the CAFs were also reversed (Figure S3A and S3B). These data indicated that the iCAF phenotype triggered by metabolic stress is reversible, suggesting that this process could represent a previously unappreciated aspect of CAF plasticity.

### iCAF markers are induced to a similar or greater extent by metabolic stress relative to extrinsic IL-1α stimulation

Previous studies have shown that IL-1α secreted by PDAC cells plays a major role in the induction of the iCAF phenotype^20,22,33^. To investigate how the intrinsic metabolic stress-induced inflammatory phenotype in CAFs compares to that induced by tumor cell-derived IL-1α, we treated CAFs with levels of IL-1α that are equivalent to what is found in PDAC cell-conditioned medium^22^ and compared the extent of expression of several iCAF markers relative to low glutamine with EIPA. We confirmed that IL-1α induced the expression of IL6, CCL2, CXCL1, CXCL2 and IL1A itself; however, we noted that the metabolic stress caused by low glutamine and macropinocytosis inhibition induced a broader panel of markers, and in many cases at higher levels (Figure S3C). Taken together, these data demonstrate that the intrinsic iCAF program regulated by metabolic stress can contribute to CAF subtype identity to a similar extent as non-autonomous reprogramming.

### The metabolic stress-induced inflammatory CAF signature is driven by MEK/ERK signaling

We and others have previously demonstrated that glutamine stress activates MEK/ERK signaling in PDAC cells^26,34^. Similarly, we found that low glutamine enhanced the levels of phospho-ERK (p-ERK) in the murine CAFs (Figure 3A). Macropinocytosis inhibition also enhanced levels of p-ERK and these levels were heightened when combined with low glutamine (Figure 3A). In the patient-derived human CAFs, we found that this combination also enhanced p-ERK levels, together with the metabolic stress marker SESN2 (Figure 3B). Importantly, linking the signaling output to protein scavenging, the addition of serum albumin rescued the enhancement of metabolic stress and MEK/ERK signaling (Figure 3C). To assess the contribution of MEK/ERK signaling to the induction of the inflammatory phenotype in CAFs, we treated mouse CAFs with SCH772984, a selective ERK inhibitor (ERKi), or trametinib, a selective inhibitor of MEK (MEKi). We found that the induction of iCAF markers caused by metabolic stress was suppressed by inhibiting MEK/ERK signaling (Figure 3D). Importantly, this was not the case for inhibitors of other stress kinases, including p38i, JNK, PI3K, or AKT (Figure S4A). To further study the molecular mechanisms underlying the intrinsic inflammatory CAF phenotype, we next assessed the expression dynamics of the various inflammatory markers. We found four ‘early phase’ markers (IL1A, CCL2, CXCL1 and CXCL2) that were first induced at approximately 4 hours post-EIPA treatment and three ‘late phase’ markers (IL1B, CXCL12 and PDGFRA) that were induced at approximately 16 hours post-treatment (Figure S4B). MEK/ERK signaling is known to induce the expression of downstream transcription factors (TFs) that drive the signal outputs. Since the expression of the early phase markers may be what triggers the full induction of the intrinsic inflammatory phenotype, we next focused on the promoters of these four markers to help us identify potential TFs involved. Using an *in silico* approach facilitated by the GeneHancer database^35^, we identified 22 TFs that were common to the promoters of all four early phase markers. We then screened for changes in the expression levels of these 22 TFs in conditions of low glutamine, EIPA treatment or low glutamine combined with EIPA. We found five transcription factors including ATF3, EGR1, FOS, FOSB, and FOSL1 that were transcriptionally enhanced in the combination treatment (Figures 3E and S4C). To identify which of these TFs could be contributing downstream of MEK/ERK signaling to execute the events leading to the inflammatory phenotype, we assessed their expression levels after MEK/ERK inhibition. We found that, except for ATF3, they were all completely dependent on MEK/ERK (Figure 3F). These data demonstrate that the induction of the intrinsic inflammatory CAF phenotype relies on MEK/ERK signaling, and that multiple TFs likely govern the expression of the inflammatory markers.

**Figure 3.**
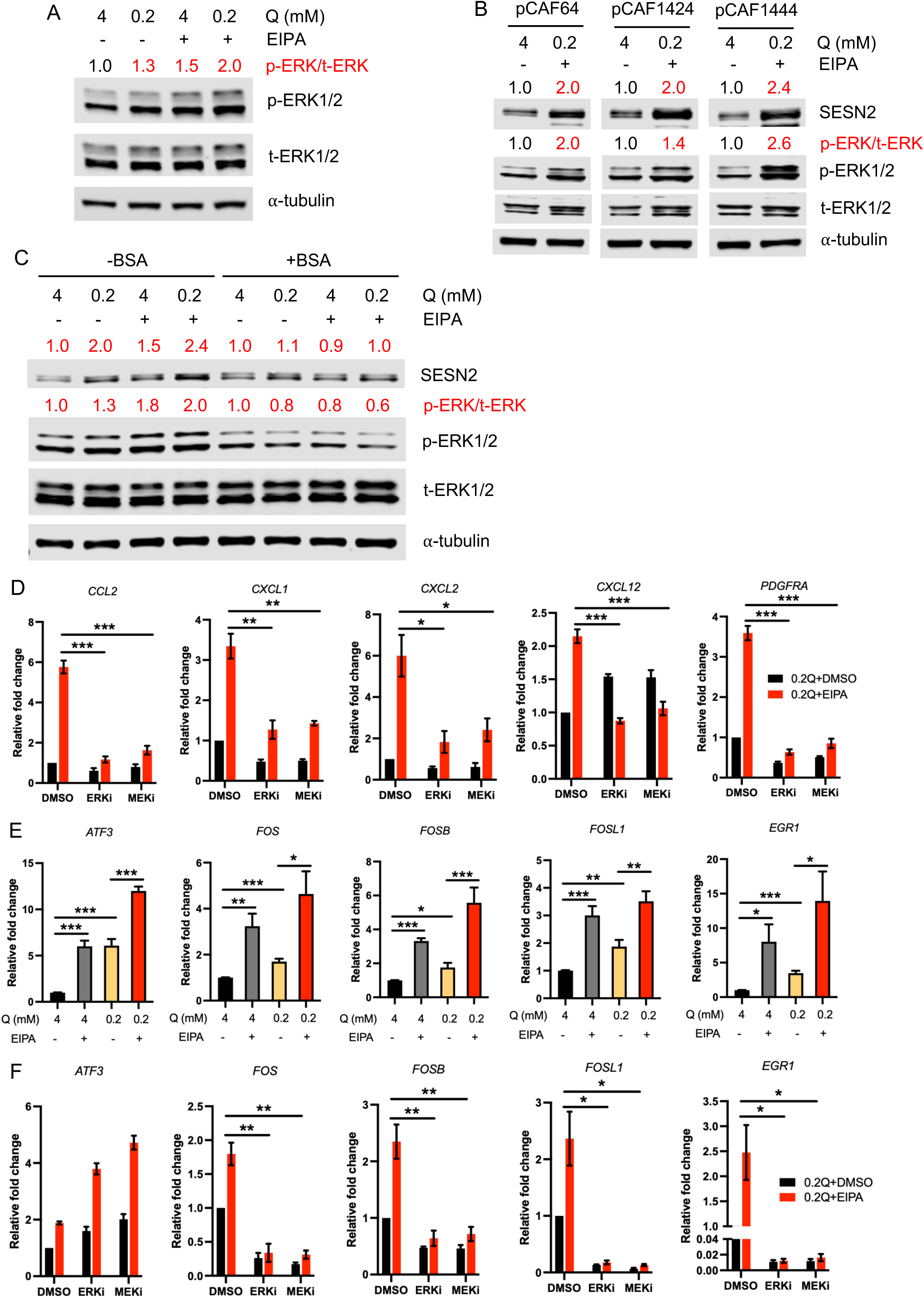
Metabolic stress-induced iCAF phenotype requires MEK/ERK signaling. (A) Mouse CAFs were treated with 4 mM or 0.2 mM Q with or without the presence of 25 μM EIPA for 4 hours. The protein levels of phospho-ERK1/2 (p-ERK1/2) and total-ERK1/2 (t-ERK1/2) were analyzed by western blots. (B) Human pCAFs were treated with 4 mM Q plus DMSO or 0.2 mM Q plus 25 μM EIPA for 24 hours. The protein levels of SESN2, p-ERK1/2 and t-ERK1/2) were analyzed by western blots. (C) Mouse CAFs were treated with 0.2 mM Q with DMSO or EIPA with or without the presence of 4% BSA for 4 hours. The protein levels of SESN2, p-ERK1/2 and t-ERK1/2) were analyzed by western blots. (D) Mouse CAFs were pre-treated with ERK inhibitor (ERKi) or MEK inhibitor (MEKi) for 1 hour and then treated with 4 mM or 0.2 mM Q with or without the presence of 25 μM EIPA for 24 hours. The mRNA levels of iCAF-related markers were analyzed by qPCR. Data are shown as mean ± SEM, unpaired t-test. *p<0.05, **p<0.01, ***p<0.001. (E) Mouse CAFs were treated with 4 mM or 0.2 mM Q with or without the presence of 25 μM EIPA for 4 hours. The mRNA levels of the transcription factors were analyzed by qPCR. Data are shown as mean ± SEM, one-way ANOVA and unpaired t-test. *p<0.05, **p<0.01, ***p<0.001. (F) Mouse CAFs were pre-treated with ERKi or MEKi for 1 hour and then treated with 4 mM or 0.2 mM Q with or without the presence of 25 μM EIPA for 6 hours. The mRNA levels of the transcription factors were analyzed by qPCR. Data are shown as mean ± SEM, unpaired t-test. *p<0.05, **p<0.01, ***p<0.001.

### Macropinocytosis supports CAF subtype identity and heterogeneity *in vivo*

The impact of macropinocytosis inhibition on tumor growth dynamics has previously only been assessed in xenograft models^28,36,37^; therefore, we tested whether EIPA administration could suppress macropinocytosis and tumor growth in a syngeneic heterotopic mouse model where murine PDAC cells (KPC cells) are implanted subcutaneously in C57BL/6 mice^38^. When the subcutaneous tumors attained a volume of ∼100 mm^3^, animals were treated with vehicle control or 10 mg/kg EIPA via intraperitoneal injection every two days for two weeks. We found that EIPA treatment significantly suppressed tumor growth and inhibited intratumoral macropinocytosis in both the tumor cells and the α-SMA^+^ CAFs (Figures S5A-S5D). To assess whether macropinocytosis inhibition induces an inflammatory phenotype in CAFs *in vivo*, we employed an orthotopic syngeneic implantation model that better recapitulates the histopathological complexity of the human disease. KPC cells were injected directly into the pancreata of C57BL/6 mice to generate orthotopic tumors and 10 days post-implantation the animals were treated with EIPA. Like the heterotopic model, we found that EIPA lessens the growth of the orthotopic tumors and inhibits macropinocytic uptake in both the tumor cells and the CAFs (Figures 4A-4E). The reduction in tumor growth upon treatment with EIPA was accompanied by a decrease in proliferation and an increase in apoptotic cell death, as assessed by anti-Ki-67 and anti-cleaved caspase 3 immunostaining, respectively (Figures S5E-S5H). To broadly assess the effects of macropinocytosis inhibition on the expression of inflammatory markers in PDAC tumors, we performed multiplex gene expression analysis using the nanoString nCounter immune profiling panel on PDAC tumors treated with vehicle or EIPA. We found that most of the iCAF-related markers and numerous other pro-inflammatory factors were increased by EIPA treatment relative to the vehicle-treated control tumors (Figures 4F and S5I). We next examined whether EIPA treatment alters CAF content and subtypes *in vivo*. We dissociated the tumors after EIPA administration and stained the CAFs with specific subtype markers as previously described^7^. We found that EIPA increased the iCAF population and concomitantly decreased the myCAF population, while apCAFs were unaffected (Figures 4G and 4H). We further validated these findings by immunohistochemical (IHC) staining. Although EIPA treatment did not alter the overall number of CAFs within the tumors, as assessed by podoplanin (PDPN) staining, a pan-CAF marker^15^ (Figure S5J and S5K), we did observe a significant decrease in α-SMA^+^ CAFs in non-peripheral regions of the tumors (Figures S5L-S5O). This is consistent with the notion that both macropinocytosis inhibition and glutamine depletion need to co-occur to efficiently trigger CAF subtype identity alterations since previous studies demonstrated that glutamine levels are most depleted in the non-peripheral regions of the PDAC tumors^26^. We further immunostained the tumors for TAGLN, a second marker of myCAFs, and found that it was also decreased within the EIPA-treated tumors, confirming a reduction of myCAFs *in vivo* (Figures 4I and 4J). Conversely, staining with anti-PDGFRα, an iCAF marker^11,15^, showed that EIPA treatment significantly increased the iCAF content within the tumor (Figures 4K and 4L). To assess whether ERK activation is associated with the CAF subtype alterations we observed in the tumors, we co-stained tumor sections with anti-p-ERK and anti-PDPN. We found that there were increased levels of p-ERK in the CAFs following EIPA treatment relative to the vehicle-treated controls (Figures 4M and 4N). In addition, we found EIPA treatment upregulated the expression of several targets of p-ERK, as assessed via the nanoString panel, further indicating the intratumoral activation of MEK/ERK signaling (Figure S5P). Overall, these results indicate that macropinocytosis controls CAF subtype heterogeneity *in vivo*, suggesting that inhibiting this uptake pathway could broadly provoke a proinflammatory tumor microenvironment.

**Figure 4.**
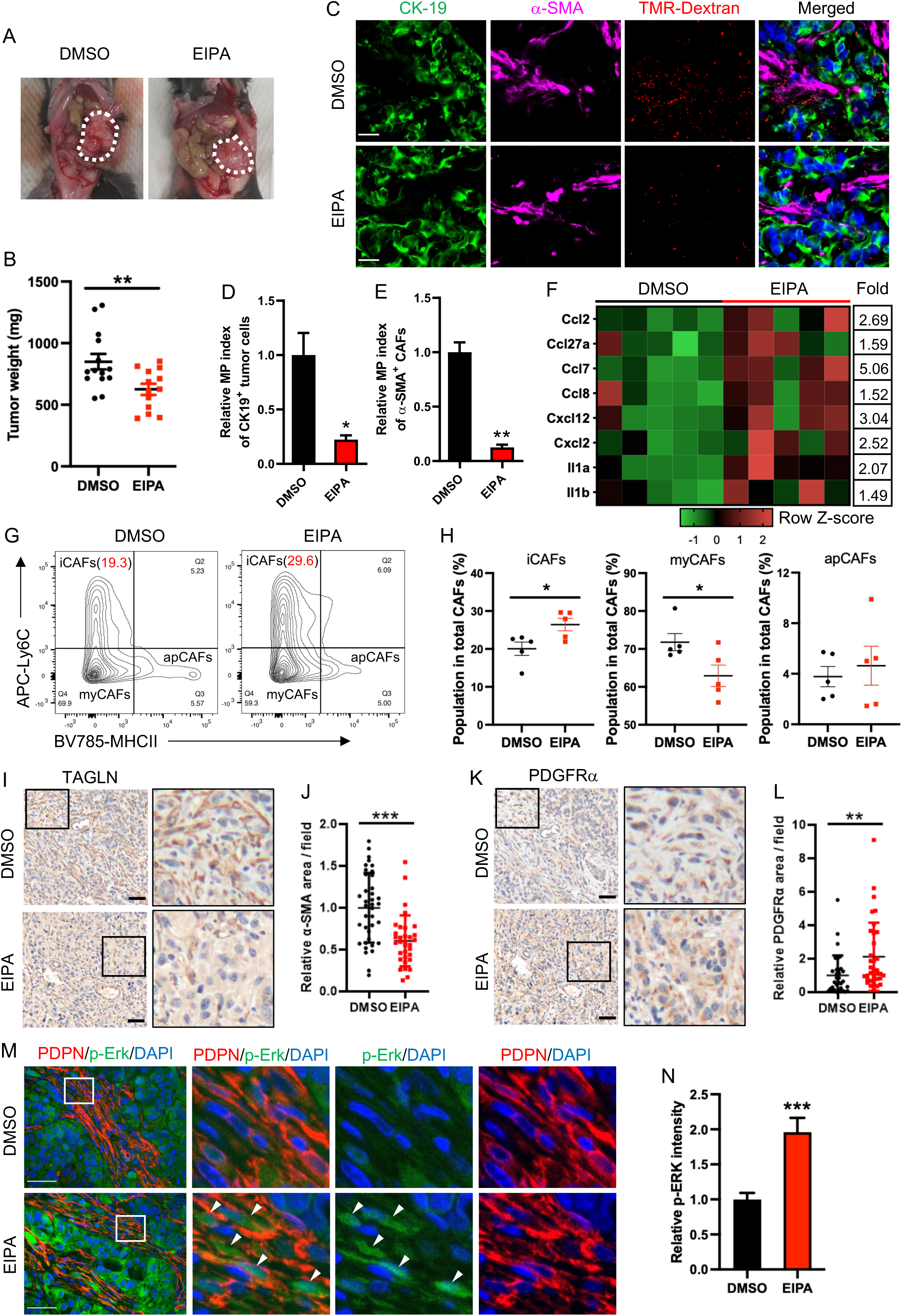
Macropinocytosis inhibition suppresses PDAC tumor growth and modulates CAF heterogeneity *in vivo*. (A and B) Representative images (A) and total tumor weights (B) of KPC orthotopic tumors treated with DMSO (vehicle, n=14) or EIPA (n=13) at 10 mg/kg for two weeks. Tumors are outlined with white dashed lines. Data are shown as mean ± SEM, unpaired t-test. *p<0.05. (C-E) Ex-vivo macropinocytic uptake of TMR-labeled dextran was performed on KPC orthotopic tumors treated with DMSO or EIPA. Tumor cells and myCAFs were labeled with CK-19 (green) or α-SMA (magenta), respectively (C). The relative macropinocytic (MP) index was calculated by the average particle intensity per area of the tumor cells (D) or the myCAFs (E), and normalized to the control. Data are shown as mean ± SEM, n_(DMSO)_=4; n_(EIPA)_=5. Unpaired t-test. *p<0.05, **p<0.01. Scale bar: 20 μm. (F) Heatmap depicting expression of iCAF-related genes in DMSO (n=5) or EIPA (n=5)-treated PDAC tumors using nanoString nCounter mouse PanCancer Immune Profiling Panel. Fold change in mRNA levels is shown for each gene. (G and H) EIPA-treated PDAC tumors were dissociated and stained with a panel of markers for CAF characterization, followed by flow cytometry analysis. From CD45^-^CD31^-^EpCAM^-^PDPN^+^ total CAFs, three CAF subtypes were identified using the markers Ly6C and MHC-II, as indicated. Representative flow cytometric results of tumors with or without EIPA treatments were shown (G). The population distribution for each CAF subtype was determined (H). Data are shown as mean ± SEM, n=5. Unpaired t-test. *p<0.05. (I and J) Representative images (I) and quantification (J) of anti-TAGLN staining in KPC orthotopic tumors treated with DMSO or EIPA. ∼35 fields (20x) from 6 tumors for each group were analyzed. Data are shown as mean ± SD, unpaired t-test. ***p<0.001. Scale bar: 50 μm. (K and L) Representative images (K) and quantification (L) of anti-PDGFRα staining in KPC orthotopic tumors treated with DMSO or EIPA. ∼35 fields (20x) from 6 tumors for each group were analyzed. Data are shown as mean ± SD, unpaired t-test. **p<0.01. Scale bar: 50 μm. (M and N) Representative images (M) and quantification (N) of anti-p-ERK1/2 staining in KPC orthotopic tumors treated with DMSO or EIPA. ∼80 fields (40x) from 5 tumors for each group were analyzed. Data are shown as mean ± SEM. Unpaired t-test. ***p<0.001. Scale bar: 50 μm.

### Macropinocytosis inhibition suppresses metastatic progression in combination with immunotherapy

In PDAC, the desmoplastic response mediated by myCAFs is thought to play a role in dampening the infiltration of T cells^39–41^. Since we observed that EIPA decreased the myCAF content in favor of a more iCAF phenotype within the tumors, we next examined whether macropinocytosis inhibition could affect the immune landscape. We performed immune profiling by flow cytometry on orthotopic KPC tumors and found that EIPA treatment increased the recruitment of CD4^+^ and CD8^+^ T cells (Figure S6A). IHC staining confirmed that macropinocytosis inhibition promotes the intratumoral infiltration of CD4^+^ and CD8^+^ T cells (Figures 5A-5D). Flow cytometry and IHC staining revealed that the orthotopic tumors did not display significant changes in the overall number of total hematopoietic cells or other immune cells (Figures S6A-S6C). Due to the highly immunosuppressive PDAC TME, intratumoral T cells display exhaustion and co-express the markers PD-1 and Tim-3^42^. Analysis of intratumoral T cells revealed that despite higher levels of infiltration, T cells still co-expressed exhaustion markers (Figure S6D). This finding suggested that the observed effects on tumor growth may not be reliant on T cells. Indeed, EIPA treatment significantly reduced the growth of KPC-derived orthotopic tumors in T cell-deficient nu/nu mice to a similar extent as was observed in C57BL/6 mice (Figure S6E). We also analyzed T cell infiltration in heterotopic PDAC tumors by IHC staining and found that tumors from animals treated with EIPA displayed significantly elevated numbers of intratumoral CD4^+^ and CD8^+^ T cells relative to controls (Figures S6F-S6I). These findings indicated that the reorganization of the stromal architecture by macropinocytosis inhibition is accompanied by enhanced CD4^+^ and CD8^+^ T cell infiltration.

**Figure 5.**
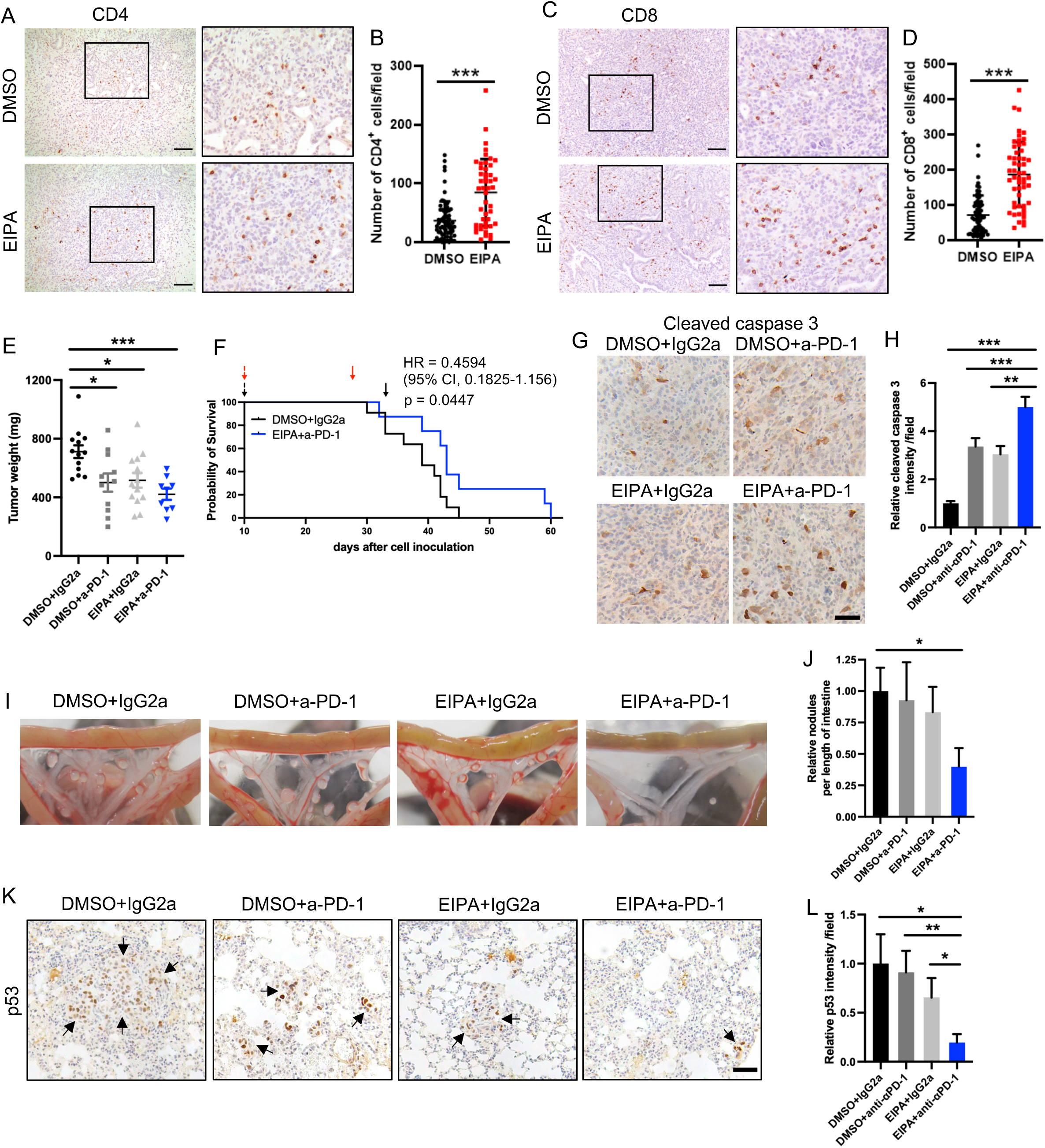
Macropinocytosis inhibition combined with immune checkpoint blockade suppresses PDAC progression. (A and B) Representative images (A) and quantification (B) of anti-CD4 staining in KPC orthotopic tumors treated with DMSO or EIPA. ∼70 fields (20x) from 6 DMSO-treated tumors and ∼50 fields (20x) from 6 EIPA-treated tumors were analyzed. Data are shown as mean ± SD, unpaired t-test. ***p<0.001. Scale bar: 100 μm. (C and D) Representative images (C) and quantification (D) of anti-CD8 staining in KPC orthotopic tumors treated with DMSO or EIPA. ∼70 fields (20x) from 6 DMSO-treated tumors and ∼50 fields (20x) from 6 EIPA-treated tumors were analyzed. Data are shown as mean ± SD, unpaired t-test. ***p<0.001. Scale bar: 100 μm. (E) Mice bearing orthotopic PDAC tumors were treated with EIPA and/or anti-PD-1 antibody. 27 days post-implantation, tumors were harvested and weighed. Data are shown as mean ± SEM; n_(DMSO+IgG2a)_=13; n_(DMSO+a-PD-1)_=11; n_(EIPA+IgG2a)_=13; n_(EIPA+a-PD-1)_=9. One-way ANOVA. *p<0.05, ***p<0.001. (F) Mice bearing orthotopic PDAC tumors were treated with EIPA+anti-PD-1 antibody or vehicle+isotype antibody. Overall survival was analyzed using Kaplan-Meier curves with the log-rank test. n_(DMSO+IgG2a)_=11; n_(EIPA+a-PD-1)_=8. (G and H) Representative images (G) and quantification (H) of anti-cleaved caspase-3 staining in KPC orthotopic tumors after treatment. Images were analyzed: DMSO+IgG2a, 54 fields (20x) from 4 tumors; EIPA+IgG2a, 39 fields (20x) from 5 tumors; DMSO+a-PD-1, 43 fields (20x) from 5 tumors; EIPA+a-PD-1, 32 fields (20x) from 4 tumors. Data are shown as mean ± SD, One-way ANOVA. **p<0.01, ***p<0.001. Scale bar: 50 μm. (I and J) Representative images of intestinal macrometastases (I), and the quantification of average number of nodules per length of intestine (J). Unpaired t-test. *p<0.05. (K and L) Representative images (K) and quantification (L) of anti-p53 staining in the lungs. 25 fields (20x) from 5 lungs for each group were analyzed. Data are shown as mean ± SD, One-way ANOVA. ***p<0.001. Scale bar: 50 μm.

Previous studies have shown that immunotherapy in PDAC tumors that display enhanced T cell infiltration has little benefit on the growth of the primary tumors but does significantly reduce metastasis^43^. Despite the T cells maintaining their exhaustion markers within the PDAC tumors treated with EIPA, we thought that there could be therapeutic potential in combining macropinocytosis inhibition with immune checkpoint blockade (ICB), especially from the perspective of metastatic progression. To test this idea, we co-treated orthotopic PDAC tumor-bearing mice with EIPA and an anti-PD-1 antibody (Figure S7A). We observed modest effects on the growth of the primary tumors and overall survival with the combination therapy (Figures 5E, 5F, S7B). The combination therapy significantly enhanced apoptosis and CD8^+^ T cell infiltration within the tumors (Figures 5G, 5H, S7C and S7D). We next sought to examine the effects of the combination therapy on metastasis to various organs, including the intestine, liver, diaphragm, and lung. Of note, we found that relative to the control, the EIPA and anti-PD-1 combination therapy significantly decreased the incidence of metastasis to any organ site (Figures S7E and S7F). Treatment with anti-PD-1 alone had no effect, while EIPA alone moderately reduced the overall incidence of macrometastases (Figures S7E and S7F). For specific organs, we found that the combination therapy reduced the incidence of macrometastases to the liver, intestine, and diaphragm relative to the control (Figure S7E and S7F). To specifically assess metastatic tumor burden, we further quantified the number of metastatic nodules per length of intestine and found that the combination therapy significantly reduced nodule number (Figures 5I and 5J). For the lung tissue, we evaluated micrometastases by H&E staining and detected infiltrating PDAC cells in the lung by IHC staining with anti-p53. We found that the combination therapy significantly decreased the number of micrometastases to the lung (Figures 5K, 5L and S7G). Overall, our data indicate that the inhibition of macropinocytosis can suppress PDAC progression in combination with immunotherapy.

### Drug delivery and chemotherapeutic responses are enhanced by macropinocytosis inhibition

In accordance with macropinocytosis inhibition decreasing myCAF content within the tumors, we found that collagen deposition in EIPA-treated orthotopic tumors was decreased relative to the vehicle-treated controls (Figures 6A and 6B). The extracellular matrix proteins (ECMs) secreted by CAFs have been demonstrated to cause blood vessel compression and collapse within PDAC tumors, and depletion of the ECMs has been shown to drive re-expansion of the tumor vasculature and to enhance drug delivery to the tumors^44–46^. Interestingly, IHC staining with anti-CD31, which labels the tumor vasculature, revealed that the blood vessels in both heterotopic and orthotopic PDAC tumors were significantly expanded by EIPA (Figures 6C, S8A and S8B). We surmised that this vascular expansion caused by EIPA administration could improve drug delivery to the tumors. To test this idea, we exploited the intrinsic auto-fluorescence of doxorubicin (Dox), a small molecule chemotherapeutic drug that is used as a therapy in several different cancers. We treated orthotopic tumor-bearing animals with Dox infusion following EIPA treatment. We found that EIPA significantly increased Dox delivery to the tumors as visualized by fluorescent accumulation of Dox in the cell nuclei relative to the vehicle-treated control tumors (Figures 6D, 6E, S8C and S8D). These data suggested that EIPA could be leveraged to enhance the delivery of chemotherapeutic drugs to the PDAC tumors. To test this idea, we next assessed the anti-tumor activity of EIPA as a pre-treatment followed by the administration of gemcitabine, a first-line chemotherapeutic drug given to patients for the treatment of advanced PDAC. We treated orthotopic PDAC tumor-bearing mice with five doses of EIPA followed by a single dose of gemcitabine (Figure 6F). Under these treatment conditions, neither EIPA nor gemcitabine significantly impacted tumor growth (Figure 6G). Importantly, EIPA treatment followed by gemcitabine significantly and synergistically suppressed tumor growth as determined by calculating the combination index (CI)^47^ (Figure 6G). The reductions in tumor growth were accompanied by decreased cell proliferation and increased apoptosis in the tumors (Figures 6H, 6I, S8E and S8F). We also observed that pre-treatment with EIPA followed by gemcitabine suppressed lung micrometastases (Figures 6J and 6K). Altogether, our data indicate that the effects of macropinocytosis inhibition on the stromal architecture can be harnessed to enhance intratumoral delivery of chemotherapeutic drugs and improve their effectiveness in PDAC.

**Figure 6.**
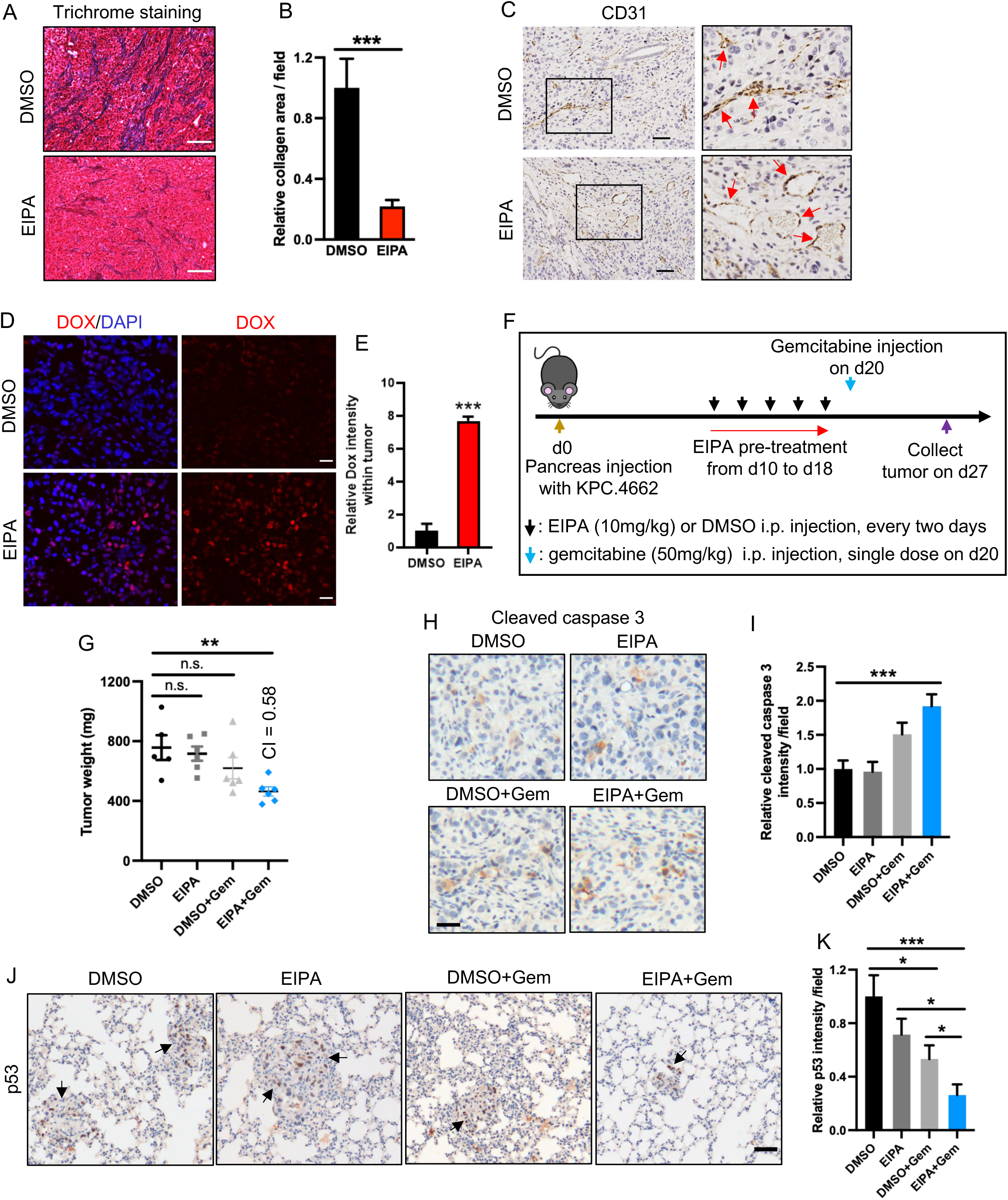
Macropinocytosis inhibition enhances drug delivery and therapeutic response to gemcitabine. (A and B) Representative images (A) and quantification (B) of collagen via trichrome staining in KPC orthotopic tumors treated with DMSO or EIPA. 28 fields (10x) from 5 DMSO-treated tumors and 25 fields (10x) from 5 EIPA-treated tumors were analyzed. Data are shown as mean ± SD, unpaired t-test. ***p<0.001. Scale bar: 100 μm. (C) Representative images of anti-CD31 staining in KPC orthotopic tumors treated with DMSO or EIPA. Red arrows indicate CD31-labeled blood vessels. Scale bar: 50 μm. (D and E) Representative images (B) and quantification (C) of doxorubicin (DOX) delivery in KPC orthotopic tumors treated with DMSO or EIPA. Nuclei were co-stained with DAPI. Data are shown as mean ± SEM (n=3). Unpaired t-test. ***p<0.001. Scale bar: 20 μm. (F) Schematic depicting the treatment protocol for EIPA pre-treatment and gemcitabine administration. (G) Mice with orthotopic PDAC tumors were treated with EIPA and/or gemcitabine. 27 days post-implantation, tumors were harvested and weighed. Data are shown as mean ± SEM; n_(DMSO)_=5; n_(EIPA)_=6; n_(DMSO+Gem)_=6; n_(EIPA+Gem)_=6. One-way ANOVA. **p<0.01, n.s., not significant. CI: combination index. (H and I) Representative images (H) and quantification (I) of anti-cleaved caspase-3 staining in KPC orthotopic tumors after treatment. Images were analyzed: DMSO, 56 fields (20x) from 5 tumors; EIPA, 67 fields (20x) from 6 tumors; DMSO+Gem, 57 fields (20x) from 5 tumors; EIPA+Gem, 70 fields (20x) from 6 tumors. Data are shown as mean ± SD, One-way ANOVA. ***p<0.001, n.s., not significant. Scale bar: 30 μm. (J and K) Representative images (J) and quantification (K) of anti-p53 staining in lungs. 25-30 fields (20x) for each group were analyzed. Data are shown as mean ± SD, One-way ANOVA. *p<0.05, ***p<0.001, n.s., not significant. Scale bar: 50 μm.

## Discussion

CAFs are a crucial component of the PDAC stroma and have been reported to be heterogeneous by lineage, phenotype and function^14,15,17,20^. Numerous CAF subtypes have been identified in PDAC tumors, including myCAFs, iCAFs, and apCAFs; and CAF plasticity has been reported to be regulated by secreted factors, such as IL-1/TGFβ and hedgehog ligands. Biffi *et al*. established that IL1 and TGFβ are two extrinsic signaling molecules to drive myCAF or iCAF phenotype, respectively, through regulating JAK/STAT signal pathway^20^. Steele *et al*. found that the inhibition of hedgehog signaling using LDE225, a SMO inhibitor, significantly reduces myCAFs but expands iCAF population^21^. What’s more, Mello *et al*., Schworer *et al*., and Singh *et al*. demonstrated that IL1-dependent iCAF phenotype could be potentiated by hypoxia signaling, or extrinsic p38 MAPK pathway, respectively^22,33^. In our study, we have demonstrated for the first time that macropinocytosis is dialed up in CAFs as an adaptive metabolic pathway to maintain CAF subtype identity under glutamine stress. Importantly, we show that macropinocytosis inhibition, when combined with glutamine depletion, induces an intrinsic iCAF phenotype while reducing myCAF properties. This is physiologically relevant to PDAC because human patient specimens, as well as multiple murine tumor models, have revealed that these tumors are glutamine deficient^25–27^, leading to intratumoral metabolic stress^25,34^. Macropinocytosis as an important protein-scavenging pathway has been found to be utilized by both cancer cells and CAFs to circumvent metabolic stress, supporting their fitness and function^7,28,48,49^. In addition, it has been demonstrated that the adaptive induction of macropinocytosis occurs predominantly in the cores of PDAC tumors that display enhanced glutamine depletion relative to the tumor periphery^26^. Consistent with these findings, in this study, we observed significant reduction of myCAFs in the central intratumoral regions of the PDAC tumors upon macropinocytosis inhibition, while their prevalence in the peripheral regions was not affected. These data indicate that the induction of macropinocytosis in CAFs spatially correlates with the level of glutamine depletion *in vivo* and functions to maintain their subtype identity. Activation of the MEK/ERK pathway has been shown to be elevated in tumor cells in response to glutamine depletion-associated metabolic stress^25,34^. In this study, we demonstrated that metabolic stress-induced intrinsic iCAF phenotype requires the activation of MEK/ERK signaling, likely due to ERK-dependent expression of transcription factors driving early-phase iCAF marker transcription.

The majority of PDAC tumors are immunologically “cold” tumors with poor T cell infiltration and a highly immunosuppressive microenvironment. In several tumor models, a dense extracellular matrix (ECM) has been shown to hinder T cell infiltration, acting as a physical barrier^50–52^. Previous research has demonstrated that reducing collagen density can promote T cell infiltration into PDAC tumors^53,54^. In this study, we have demonstrated that inhibiting macropinocytosis leads to a CAF subtype shift from myCAFs to iCAFs, resulting in decreased collagen deposition and increased CD8^+^ and CD4^+^ T cell infiltration. Early studies in genetically engineered mouse models revealed that a significant depletion of α-SMA^+^ CAFs decreased collagen deposition and tumor fibrosis but also lowered the overall infiltration of CD45^+^ immune cells and the effective CD8^+^/Treg ratio, indicating the diverse functions of α-SMA^+^ CAFs beyond physical barriers^19,55^. A recent study demonstrated that LDE225, a novel inhibitor of hedgehog signaling, promoted a significant phenotypic switch from myCAF to iCAF populations, which increased the infiltration of a specific subset of myeloid cells and macrophages without significantly affecting other types of immune cells^21^. These conflicting results may stem from the complex heterogeneity within the myCAF and iCAF populations themselves, as revealed by single-cell RNA sequencing identifying distinct subsets such as ecm-myCAF, TGFβ-myCAF, wound-myCAF, detox-iCAF, and IL-iCAF^56^. Interestingly, they found that the abundance of ecm-myCAF and TGFβ-myCAF was associated with an immunosuppressive environment enriched in Tregs, while, in contrast, the abundance of detox-iCAF and IL-iCAF was correlated with CD8^+^ T cell infiltration^56^. Our study demonstrates that metabolic stress specifically induces the expression of many iCAF and iCAF-related inflammatory markers, such as T cell-attractive chemokines CCL7, CCL8 and CCL27, compared to IL1 stimulation. In the future, it could be useful to understand CAF heterogeneity and plasticity more comprehensively within these diverse subsets of myCAF and iCAF populations in the context of metabolic stress. Importantly, in our study, we found that simply enhancing CD8^+^ T cell infiltration does not seem to significantly impact the growth of the primary PDAC tumors upon treatment with immunotherapy. We did, however, observe that metastasis was suppressed. These findings are consistent with a previous study by Li *et al*. that demonstrated that immunotherapy in the setting of tumor cell-defined T cell infiltration did not correlate with PDAC tumor growth but did restrict tumor metastasis.

The predominant CAF subtype found within the PDAC stroma is myCAFs, which are known for their substantial deposition of ECM proteins that contribute to desmoplasia and drive PDAC tumor progression. CAFs exhibit rewired metabolism to sustain their pro-tumorigenic phenotype^57^. Eckert *et al*. identified Nicotinamide N-methyltransferase (NNMT) as a key epigenetic regulator in CAFs to reduce histone methylation and increase expression of CAF markers, cytokines, and ECM proteins^58^. Also, Kay *et al*. found that pyrroline-5-carboxylate reductase 1 (PYCR1) is upregulated in CAFs to promote glutamine-derived proline synthesis and collagen deposition^59^.

In our study, we demonstrate a novel role for macropinocytosis in maintaining the myCAF phenotype under glutamine stress, facilitating ECM secretion and PDAC progression. ECM-caused desmoplasia has been shown to increase the interstitial fluid pressure and compress the tumor vasculature in PDAC, which leads to poor drug perfusion^60,61^. Olive *et al*. depleted myCAFs in Pdx1^Cre^;Kras^LSL-G12D/+^;p53^R172H/+^ (PKC) tumors through systemic administration of IPI-926, a Sonic hedgehog (Shh) inhibitor, demonstrated a reduction in ECM deposition and an increase in intratumoral vascular density and gemcitabine delivery, leading to transient stabilization of disease^62^. However, there were some opposing studies showing varying outcomes of depleting the stroma with respect to tumor vasculature and drug delivery^19,55^. Contrary to stroma-depleting strategies, our approach of macropinocytosis inhibition targets cellular fitness in both tumor cells and CAFs simultaneously. We found that the tumor vasculature that is normally collapsed in our orthotopic tumors expands with macropinocytosis inhibition, enhancing drug delivery. Gemcitabine preceded by macropinocytosis inhibition suppressed PDAC tumor growth and significantly suppressed lung metastasis. Such stromal effects are more akin to those observed with treatment of calcipotriol, a vitamin D receptor ligand, which reprogrammed the CAFs to be quiescent and reduced fibrosis in PDAC tumors, resulting in increased intratumoral gemcitabine delivery and improved chemotherapy response^46^. Irrespective of whether the stroma is reprogrammed or depleted, such approaches may have therapeutic utility for drug delivery in the clinic^63–65^. Altogether, these studies indicate the promising therapeutic potential that stromal reprogramming may have in PDAC. Moreover, our work underscores that therapeutic intervention strategies that dual-modulate the tumor cells and the stroma, such as macropinocytosis inhibition, should be further explored in PDAC.

## Acknowledgments

We are grateful to members of the Commisso laboratory for their helpful comments and discussions. This work was supported by NIH grants R01CA254806 and R01CA207189 to C.C. We thank Dr. Robert Vonderheide for KPC.4662 cells. We thank Guillermina Garcia and Monica Sevilla as well as the Sanford Burnham Prebys Histology Core for performing trichrome staining. We thank Buddy Charbono and Adriana Charbono as well as the Sanford Burnham Prebys Animal facility for providing technical assistance. Sanford Burnham Prebys core services are supported by NCI Cancer Center Support grant P30CA030199.

## Author contributions

Y.Z. and C.C. conceived the study, developed methodology, validated, and analyzed the data, and wrote the original draft of the manuscript. Y.Z., L.L., S.M., J.H., C.M.G., F.C., and K.DP. performed experiments. A.M.L. and A.B. provided tissues and cells. D.A.S. performed the polar metabolomics. J.H. and L.B. supervised the flow cytometry experiments. A.M.L., J.H. and L.B. reviewed the manuscript. C.C. acquired funding for the study.

## Materials and methods

### Murine CAF isolation and characterization

Murine CAFs were isolated from Kras^LSL-G12D/+^;Trp53^LSL-R172H/+^;Pdx1-Cre;ROSA26^LSL-EYFP^ (EKPC) mice^66^. Briefly, EKPC tumor portions were collected in a sterile hood and minced with razor blades to ∼2 mm pieces. After rinse with DPBS, multiple tumor pieces were seeded in a 10 cm plate with 5 mL DMEM complete medium (high glucose, L-glutamine DMEM supplemented with 10% FBS, 100 U/mL penicillin-streptomycin (pen-strep), and 20 mM HEPES) plus 100 μg/ml gentamycin and incubated at 37 °C. Three days later, tumor pieces were removed. Spindle-shaped fibroblast clusters distant from tumor cell clusters were digested in situ with 0.25% trypsin and collected into a 24-well plate for growth. Finally, primary CAFs were immortalized with SV40 T antigen retrovirus as previously described^7^. Isolated murine CAFs were characterized by the expression of CAF markers using western blots and flow cytometry.

### Cell culture and treatments

LSL-Kras^G12D/+^; Trp53^fl/+^; Pdx1-Cre (KPC) mice-derived PDAC cells (KPC.4662) were provided by Dr. Robert Vonderheide (University of Pennsylvania). EKPC cells, KPC.4662 cells, and isolated murine CAFs were cultured in DMEM complete medium. Patients-derived CAFs were cultured in DMEM complete medium with an additional supplement of 7% FBS as previously described^7^. All the cells were cultured under 5% CO2 at 37 °C and routinely tested for mycoplasma contamination using ABM’s PCR Mycoplasma Detection Kit.

For treatments, murine CAFs were cultured in low glutamine DMEM medium containing 0.2 mM glutamine, 10% dialyzed FBS, 100 U/mL pen-strep, and 20 mM HEPES. The control DMEM medium contains 4 mM L-glutamine. 5-(N-ethyl-N-isopropyl)-amiloride (EIPA, Sigma), an inhibitor of Na+/H+ exchange, was administered to cells at a concentration of 25 μM. STO-609 (Santa Cruz), a selective inhibitor of CaMKK2, was administered to CAFs at a concentration of 60 μM. EHT 1864 (Cayman Chemical), an inhibitor of Rac subfamily of Rho GTPase, was administered to CAFs at a concentration of 20 μM. Murine IL-1α (R&D) was used at a concentration of 25 pg/mL for 32 hours. For kinase inhibition, 1 μM SCH772984 (Selleckchem), 1 μM Trametinib (Selleckchem), 10 μM SB202190 (Selleckchem), 10 μM LY294002 (CST), 2 μM MK-2206 (MedChemExpress) and 2 μM JNK-IN-8 (Selleckchem) were utilized to inhibit ERK, MEK, p38, PI3K, Akt and JNK kinases, respectively. For amino acid-stress rescue experiments, growth media was supplemented with 40 mg/mL bovine serum albumin (BSA, Millipore Sigma) fraction V for 24 hours.

### Macropinocytosis assay

*In vitro* macropinocytosis assay was performed in triplicates for each condition as previously described^67,68^. Briefly, CAFs were plated onto glass coverslips in 24-well plates overnight and then treated in 0.2 mM glutamine DMEM medium with or without EIPA, STO-609, or EHT 1864 for 24 hours. For BSA rescue experiments, 40 mg/mL BSA was added to growth media with vehicle or EIPA. Macropinocytosis was assessed following 30 min incubation at 37 °C with 1 mg/mL 70-kDa FITC-dextran (Invitrogen), followed by five times rinse with ice-cold PBS and then fixed in 3.7% formaldehyde. After nuclear staining with DAPI, coverslips were mounted onto glass slides using DAKO mounting medium (Agilent Tech). Fluorescent images were automatically captured at 40x magnification using Cytation 5 Imaging Multi-Mode Reader (BioTek). The quantification of macropinosome particles was analyzed as previously described^69^. The macropinocytic index (MP index) was calculated by the total particle intensity or area per cell and normalized to the control. At least 1500 cells were analyzed per condition.

*Ex vivo* macropinocytosis assay was performed as previously described^70^. Briefly, ∼1 mm thick cross-sections were immediately cut from freshly harvested tumors and injected at multiple sites with 20 mg/mL 10-kDa TMR-dextran (Invitrogen), followed by immersion in 400 μL TMR-dextran for 15 min. After three times of quick rinse with PBS, cross-sections were embedded in O.C.T. compound and snap-frozen on dry-ice. Tumor cells were labeled with rat anti-CK19 antibody (DSHB, 1:400), and myCAFs were labeled with rabbit anti-α-SMA antibody (CST, 1:500). Afterwards, tumor tissues were incubated with Alexa Fluor 488-conjugated goat anti-rat and Alexa Fluor 647-conjugated goat anti-rabbit secondary antibodies, followed by nuclear staining with DAPI and mounted onto glass slides with DAKO mounting medium. Images were captured at 40x magnification using the EVOS FL Cell Imaging System and analyzed using ImageJ software as previously described^70^. At least 10 fields per each tumor were captured and analyzed. The MP index was calculated by the total particle intensity per the area of tumor cells or CAFs and normalized to the control.

### Western blotting

Cells were lysed in RIPA buffer (10 mM Tris-HCl [pH 8.0], 150 mM NaCl, 1% sodium deoxycholate, 0.1% SDS, 1% Triton X-100) supplemented with 1x protease inhibitor cocktail and 1x PhosSTOP. After clearance with centrifugation at 18,000 g for 15 min at 4 °C, protein concentrations were determined by DC Protein Assay (Bio-Rad). Protein samples were separated by SDS-PAGE electrophoresis and then transferred to nitrocellulose membranes using Trans-Blot Turbo Transfer System (Bio-Rad), followed by incubation with primary antibodies: rabbit α-SMA (CST, 1:1000), rabbit E-cadherin (CST, 1:1000), rabbit SESN2 (Proteintech, 1:500), rabbit phospho-ERK1/2 (CST, 1:1000), rabbit ERK1/2 (CST, 1:1000), and mouse α-tubulin (Sigma, 1:4000). After staining with IRDye 680RD or 800CW secondary antibody (LI-COR), immunoblots were detected using Odyssey CLx imager (LI-COR).

### Quantitative real-time PCR (qRT-PCR)

Total RNA was isolated from cells using the PureLink RNA Mini Kit (Invitrogen) according to the manufacturer’s instructions. cDNA was synthesized from 1 μg total RNA using the High-Capacity cDNA Reverse Transcription Kit (Applied Biosystems). qRT-PCR was then performed using SYBR Premix Ex Taq II master mix (Takara) on the LightCycler 96 Instrument (Roche) or CFX384 touch real-time PCR system (Bio-Rad). The relative mRNA levels of target genes were calculated and normalized to the mRNA level of reference gene RPL13 with respect to Cq values. The sequences of primers used for qRT-PCR in this study are listed in the Key Resources Table.

### F-actin staining in CAFs

For F-actin staining, murine CAFs were permeabilized with 0.1% Triton X-100 and stained with 100 nM Acti-stain™ 488 phalloidin (Cytoskeleton, Inc.) for 30 min. Nuclei were stained with DAPI, followed by mounting with DAKO medium.

### Flow cytometric characterization of CAF subtypes

Isolated murine CAFs in suspension were pre-incubated with anti-CD16/32 antibody at 4 °C for 15 min and then stained with CoraLite^®^Plus (CL) 488-conjugated anti-FSP-1 (Proteintech, Cat# CL488-16105) and APC/Cy7-conjugated anti-PDPN (BioLegend, clone 8.1.1) antibodies at 4 °C for 30 min, followed by flow cytometric analysis. Flow cytometric characterization of CAF subtypes *in vivo* was performed as previously described^7^ with slight modification. Briefly, PDAC tumors were dissociated using a Tumor Dissociation Kit (Miltenyi Biotec). After purification, cells in suspension were blocked with anti-mouse CD16/32 antibody (BioLegend) at 4 °C for 15 min and then stained with the following antibodies at 4 °C for 30 minutes: Alexa Fluor 594-CD45 (BioLegend, clone 30-F11), Brilliant Violet 421-CD31 (BioLegend, clone 390), Alexa Fluor 488-EpCAM (BioLegend, clone 9C4), APC/Cy7-PDPN, Alexa Fluor 647-Ly6C (BioLegend, clone HK1.4), and Brilliant Violet 785-I-A/I-E (MHC-II) (BioLegend, clone M5/114.15.2), followed by flow cytometric analysis.

### Animal experiments

Six-week-old C57BL/6 mice or athymic nude mice (Foxn1nu/Foxn1nu) were purchased from Jackson Laboratories or Charles River, respectively, and allowed to acclimatize for at least 4 days. Mouse studies were conducted at the animal facility of Sanford Burnham Prebys Medical Discovery Institute in accordance with the mouse handling and experimental protocols that were approved by the Institutional Animal Care and Use Committee (IACUC). This study followed all the relevant ethical standards.

For heterotopic syngeneic mouse models, 2.5×10.^5^ KPC.4662 cells suspended in PBS with 50% matrigel (Corning) were injected subcutaneously into the flanks of C57BL/6 mice. Tumor sizes were measured using a digital caliper and calculated according to the formula: 1/2 x length (mm) x width (mm)^2^. When tumor sizes reach ∼100 mm^3^, mice were randomly grouped and treated with vehicle or 10 mg/kg EIPA intraperitonially (i.p.) every two days for two weeks. Tumor sizes were monitored every two days. After treatment, mice were sacrificed, and tumors were harvested for *ex vivo* macropinocytosis assay, or fixed in 10% formalin for immunohistochemistry.

For orthotopic tumor models, 2.5×10.^4^ KPC.4662 cells suspended in PBS with 50% matrigel (Corning) were injected into the pancreata of C57BL/6 or athymic nude mice. 10 days later, mice were randomly grouped and treated with vehicle or 10 mg/kg EIPA i.p. every two days for two weeks. After treatment, tumors were harvested, and tumor weights were measured. Tumor tissues were collected for *ex vivo* macropinocytosis assay or fixed for immunohistochemistry. For combination immunotherapy, 10 days after implantation, mice were randomly divided into four groups and treated with vehicle or 10 mg/kg EIPA i.p. every two days and/or with i.p. administration of 200 μg IgG2a or anti-PD-1 antibody on day 10, day 13, day 16, day 19, day 22, and day 25 post inoculation. Tumor burden and metastases were determined on day 27 post-inoculation following euthanasia. For combination chemotherapy, 10 days after KPC cells were orthotopically inoculated, mice were randomly divided into four groups. Mice were treated with vehicle or 10 mg/kg EIPA i.p. every two days for 5 doses and/or with 50 mg/kg gemcitabine i.p. on day 20 post inoculation. 27 days post inoculation, tumors were harvested, and tumor weights were measured. The outliers of tumor weights were identified by Grubb’s test and removed from data analysis. The combination index (CI) for EIPA and gemcitabine co-treatment was calculated using the standard formula: 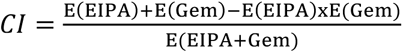 using Bliss independence approach as previously described^47^. *CI* < 1 indicates a synergistic effect between two treatments. Tumor tissues and lungs were fixed in 10% formalin for immunohistochemistry. Macrometastases in the liver, intestine, and diaphragm were visually inspected, photographically documented and recorded. For doxorubicin delivery, tumor-bearing mice were treated with vehicle or 10 mg/kg EIPA i.p. every two days for two weeks and then treated with tail vein pulse infusion of 200 μg doxorubicin. 8 min later, tumors were harvested immediately and snap frozen in O.C.T. compound for immunofluorescent staining.

### Isolation of tumor interstitial fluid (TIF) and plasma for metabolite quantification

TIF was isolated from KPC tumors as previously described^30^. Briefly, tumor-bearing mice were euthanized by cervical dislocation and tumors were rapidly dissected. After brief rinse in cold PBS and removal of extra surface liquid with filter paper, the tumors were put onto 20 μm nylon filters (Millipore Sigma) affixed atop 15 mL conical tubes and centrifuged for 10 min at 4 °C at 106 x g to collect TIF. Blood was collected from the heart of the same animals, immediately placed in EDTA-tubes (BD Vacutainer) and centrifuged at 1000 x g at 4 °C for 10 min to collect plasma. TIF and Plasma were frozen with dry ice and stored at −80 °C. Polar metabolites from 5 μL of TIF or plasma were extracted and processed for amino acids quantification by GC-MS.

### Immune profiling

C57BL/6 mice bearing orthotopic KPC.4662 tumors were implanted with an osmotic pump (Alzet, Model 1004) filled with vehicle or 20 mg/ml EIPA for drug administration over 6 days as previously described^28^. Afterwards, tumors were harvested and processed to single suspended cells as previously described^7^. Briefly, tumor portions were minced to 3-4 mm pieces and dissociated using a Tumor Dissociation Kit (Miltenyi Biotec), followed by the clearance of red blood cells with RBC lysis buffer (eBioscience) and resuspension in RPMI/2% FBS. Cell number was counted using a hemocytometer. ∼1×10.^6^ cells were stained with antibody cocktail at 4 °C for 20 min: Pacific Blue™ (PB) anti-mouse CD45.2 (Biolegend), APC anti-mouse B220 (Biolegend), FITC anti-mouse I-A^b^ (Biolegend), Brilliant Violet 605™ (BV605) anti-mouse CD8a (Biolegend), BUV395 anti-mouse CD4 (BD ebioscience), PE anti-mouse CD366 (Tim-3) (Biolegend), PE/Cy7 anti-mouse CD279 (PD-1) (Biolegend), FITC anti-mouse CD8a (Biolegend), BV605 anti-mouse NK1.1 (Biolegend), Biotin anti-mouse CD11c plus BV605 Streptavidin (Biolegend), APC anti-mouse Ly-6G (Biolegend), PE anti-mouse Ly-6C (Biolegend), APC/Cy7 anti-mouse F4/80 (Biolegend), BV480 anti-mouse CD11b (Biolegend), and anti-mouse CD16/32 antibodies. After washing twice with cold PBS/2% FBS, cells were fixed in 1% paraformaldehyde and then analyzed by flow cytometry. From each sample, 100,000 events were acquired on the BD LSRFortessa™ X-20 flow cytometer. Data analysis was performed using FlowJo v10 (BD Biosciences).

### Nanostring nCounter gene expression assay

Portions of heterotopic KPC.4662 tumors treated with vehicle or EIPA-filled osmotic pumps over 6 days were snap frozen in liquid nitrogen immediately after harvest. Frozen tumor portions were lysed on ice in the lysis buffer provided in the PureLink RNA Mini Kit (Thermo Fisher Scientific) using a Kimble pellet pestle motor, and total RNA was isolated using the kit. 50 ng RNA of each sample was hybridized using the NanoString PanCancer Immune CodeSet (Cat# XT-CSO-MIP1-12) according to the manufacturer’s instructions. Absolute read counts were quantified by the nCounter digital analyzer (nanoString).

### Doxorubicin detection in tumors

Tumor tissues with pulse infusion of doxorubicin were cut into sections with 5 μm thickness and stained with DAPI (1:2000) to label the nuclei, followed by mounting with DAKO media. Images were captured at 40x magnification using the EVOS FL Cell Imaging System and analyzed using Fiji (ImageJ) software. Doxorubicin autofluorescent images in whole tumors were acquired on a Zeiss LSM710 confocal microscope at 20X magnification. The images were taken as Z-stack under identical settings, and the maximum intensity projection images were processed using ImageJ software.

### Immunohistochemistry, trichrome and H&E staining

Formalin-fixed paraffine-embedded (FFPE) tumor tissues were cut to 5 μm sections. Immunohistochemistry staining was performed as previously described^7^. Briefly, slides with tumor sections were heated in 0.01 M citrate buffer (pH 6.0) or TE buffer (pH 9.0) for heat-induced antigen retrieval. For intracellular proteins, tumor sections were permeabilized with 0.1% Triton-X100 in TBS containing 0.1% Tween 20 (TBST) for 10 min. Afterwards, tumor sections were blocked with 10% goat or horse serum plus 1% BSA at R.T. for 1 hour and then stained with primary antibodies at 4 °C overnight, followed by incubation with biotinylated goat anti-rabbit or horse anti-goat secondary antibody (Vector Labs, BA-1000, BA-9400) at R.T. for 1.5 hours. The staining signal was developed using HRP-coupled ABC reagent (Vector Labs) and the DAB HRP Substrate Kit (Vector Labs). For p53 staining of mouse lung tissues, a mouse-on-mouse kit (Vector Labs) was used to reduce the background signaling. After nuclear counterstaining with hematoxylin, sections were dehydrated and mounted with coverslips using Permount mounting medium (Fisher Scientific). The antibodies using in this study: rabbit anti-CD4 (Abcam, 1:500), rabbit anti-CD8a (Abcam, 1:1500), rabbit anti-Ki-67 (Abcam, 1:200), rabbit anti-α-SMA (CST, 1:500), rabbit anti-cleaved caspase 3 (CST, 1:600), rabbit anti-transgelin (TAGLN, Proteintech, 1:150), goat anti-PDGFRα (R&D, 5 μg/ml), rabbit anti-CD45 (CST, 1:200), rabbit anti-CD31 (Abcam, 1:2000), and mouse anti-p53 (CST, 1:2000). For immunofluorescent staining from FFPE samples, tissue sections were stained with anti-PDPN (Thermo Fisher Scientific, 1:100) alone or co-stained with anti-phospho-ERK1/2 (CST, 1:100) and nuclei were stained with DAPI, followed by mounting with DAKO media. Trichrome and H&E staining of FFPE tumor samples were performed by Sanford Burnham Prebys Medical Discovery Institutes Histology Core Facility. Images were captured using a bright field microscope installed with INFINITY camera (Lumenera). Positive cell numbers per field and collagen area per field of trichrome staining were analyzed using Fiji (ImageJ) software.

### Statistical analysis

For all *in vitro* results, at least three independent experiments were performed, with each experiment containing two or three biological replicates derived from independent samples. For all *in vivo* results, the number of animals for each cohort is as indicated. All graphs were made using GraphPad Prism software (GraphPad). Results are shown as means; error bars represent standard error of the mean (SEM) or standard deviation (SD). Statistical significance was determined by the unpaired two-tailed Student’s t-test with or without Welch’s correction or one-way analysis of variance (ANOVA), and p values less than 0.05 were considered statistically significant (* p< 0.05, ** p< 0.01, *** p< 0.001, **** p< 0.0001).

## Supplemental Figure Legends

**Supplemental Figure 1.**
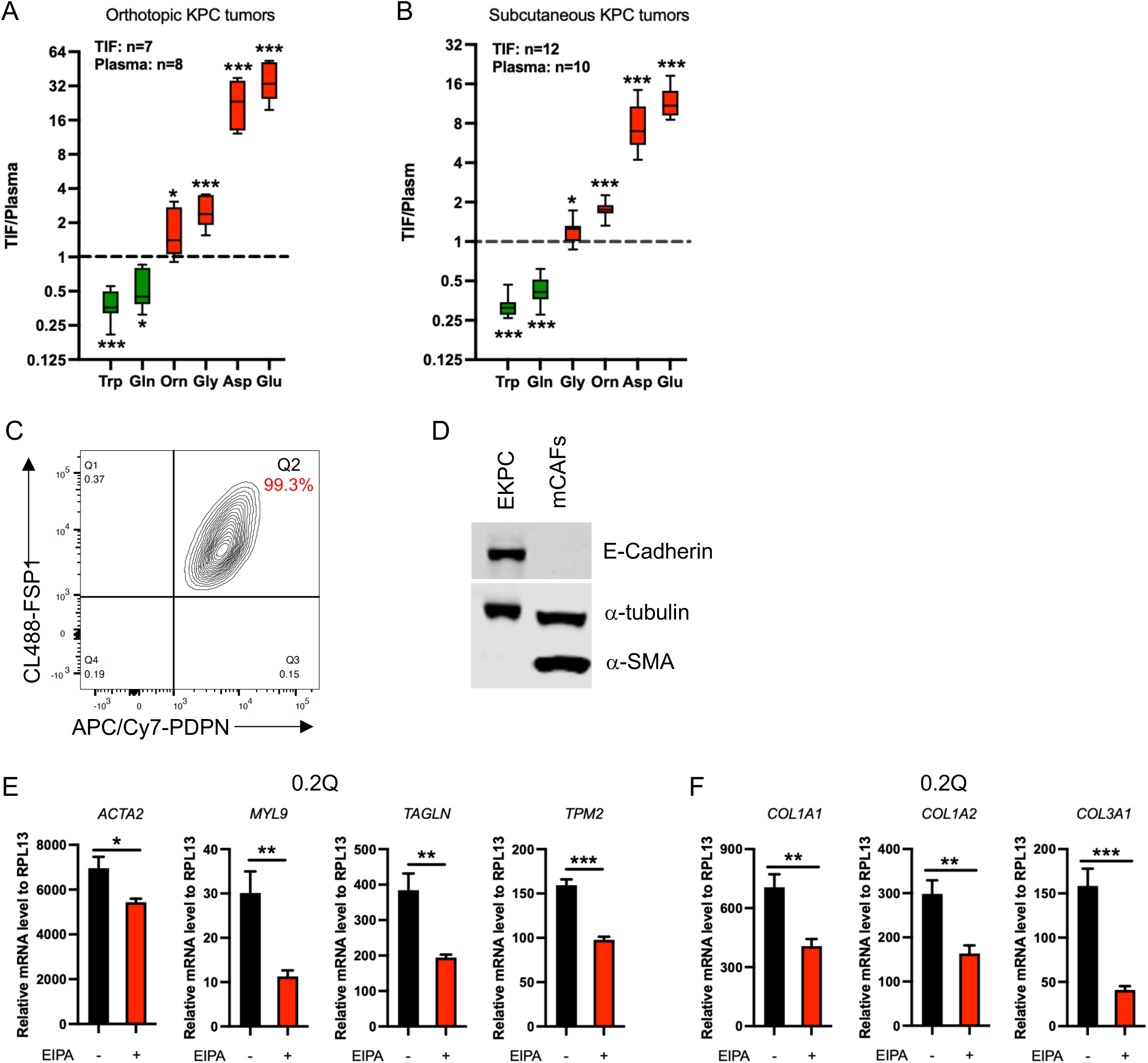
Low glutamine and macropinocytosis inhibition cooperate to induce the iCAF phenotype. (A and B) Quantitation of amino acid levels in tumor interstitial fluid (TIF) relative to the plasma in mice harboring orthotopic (A) or subcutaneous (B) KPC tumors. Data are shown as mean ± SEM, unpaired t-test. *p<0.05, ***p<0.001. (C) Characterization of CAF markers by flow cytometry. More than 99% of isolated CAFs express both podoplanin (PDPN), a pan-CAF marker, and fibroblast-specific protein 1 (FSP1). (D) Western blots of epithelial or myCAF markers in EKPC tumor cells and isolated murine CAFs. (E) mRNA expression levels of myCAF markers in CAFs treated with 0.2Q with DMSO or 25 μM EIPA for 24h. Data from at least three independent experiments are shown as mean ± SEM, unpaired t-test. *p<0.05, **p<0.01, ***p<0.001. (F) mRNA expression levels of collagen subtypes in CAFs treated with 0.2Q with DMSO or 25 μM EIPA for 24h. Data from at least three independent experiments are shown as mean ± SEM, unpaired t-test. *p<0.05, **p<0.01, ***p<0.001.

**Supplemental Figure 2.**
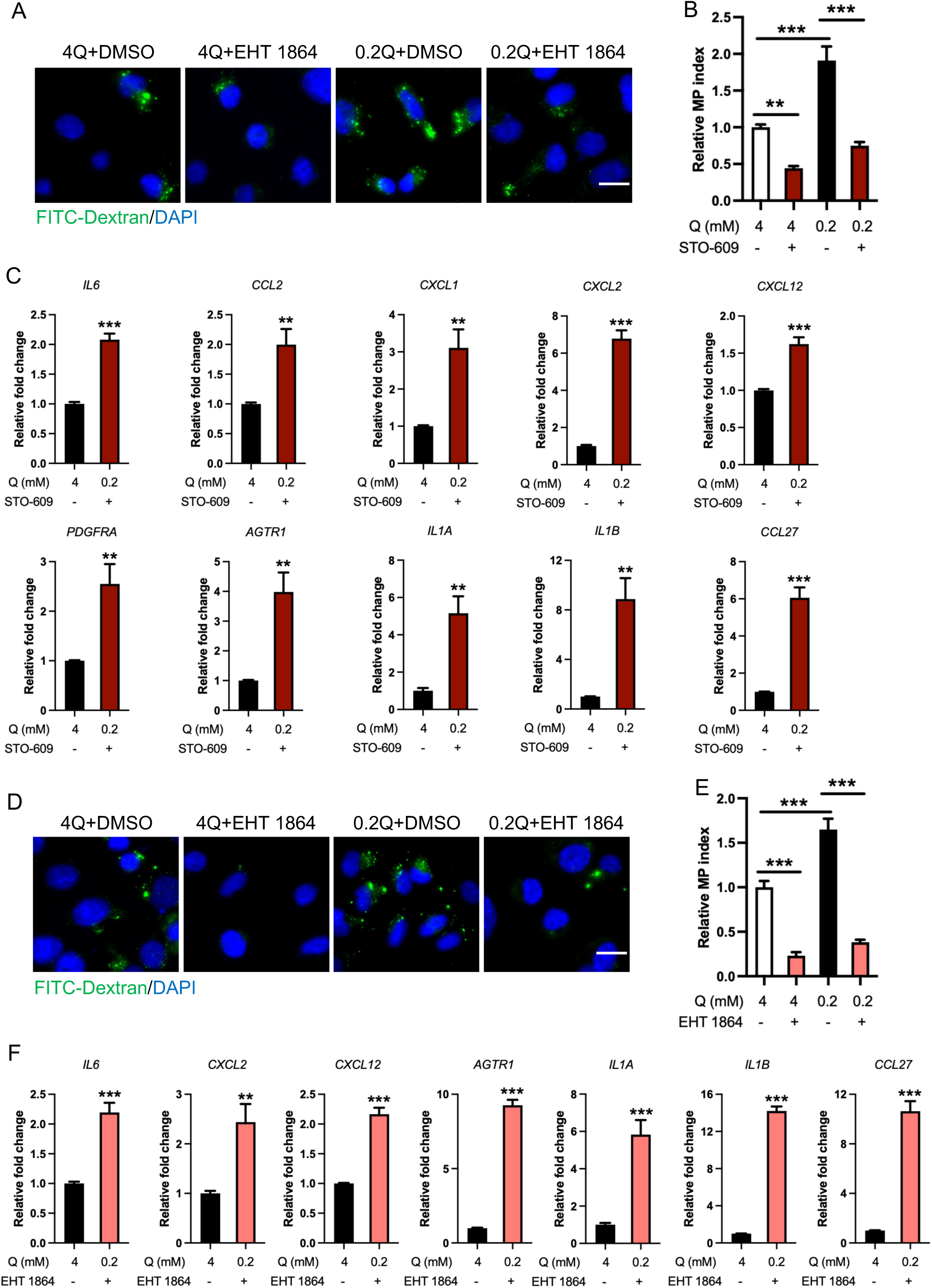
Multiple macropinocytosis inhibitors induce an intrinsic iCAF phenotype. (A and B) Representative images (A) and quantification (B) of macropinocytic uptake of FITC-labeled dextran in CAFs treated with DMSO or 60 μM STO-609 for 24 hours. Data are shown as mean ± SEM, n-3. Unpaired t-test. **p<0.01. Scale bar: 20 μm. (C) Relative mRNA expression levels of iCAF-related markers in CAFs treated with vehicle or 60 μM STO-609 for 48h, which was normalized to the vehicle-treated conditions. Data are shown as mean ± SEM, n=3. Unpaired t-test. *p<0.05, **p<0.01, ***p<0.001. (D and E) Representative images (D) and quantification (E) of macropinocytic uptake of FITC-labeled dextran in CAFs treated with DMSO or 20μM EHT 1864 for 24 hours. Data are shown as mean ± SEM, n=3. Unpaired t-test. **p<0.01. Scale bar: 20 μm. (F) Relative mRNA expression levels of iCAF-related markers in CAFs treated with vehicle or 20 μM EHT 1864 for 48 hours, which was normalized to the vehicle-treated conditions. Data are shown as mean ± SEM, n=3. Unpaired t-test. *p<0.05, **p<0.01, ***p<0.001.

**Supplemental Figure 3.**
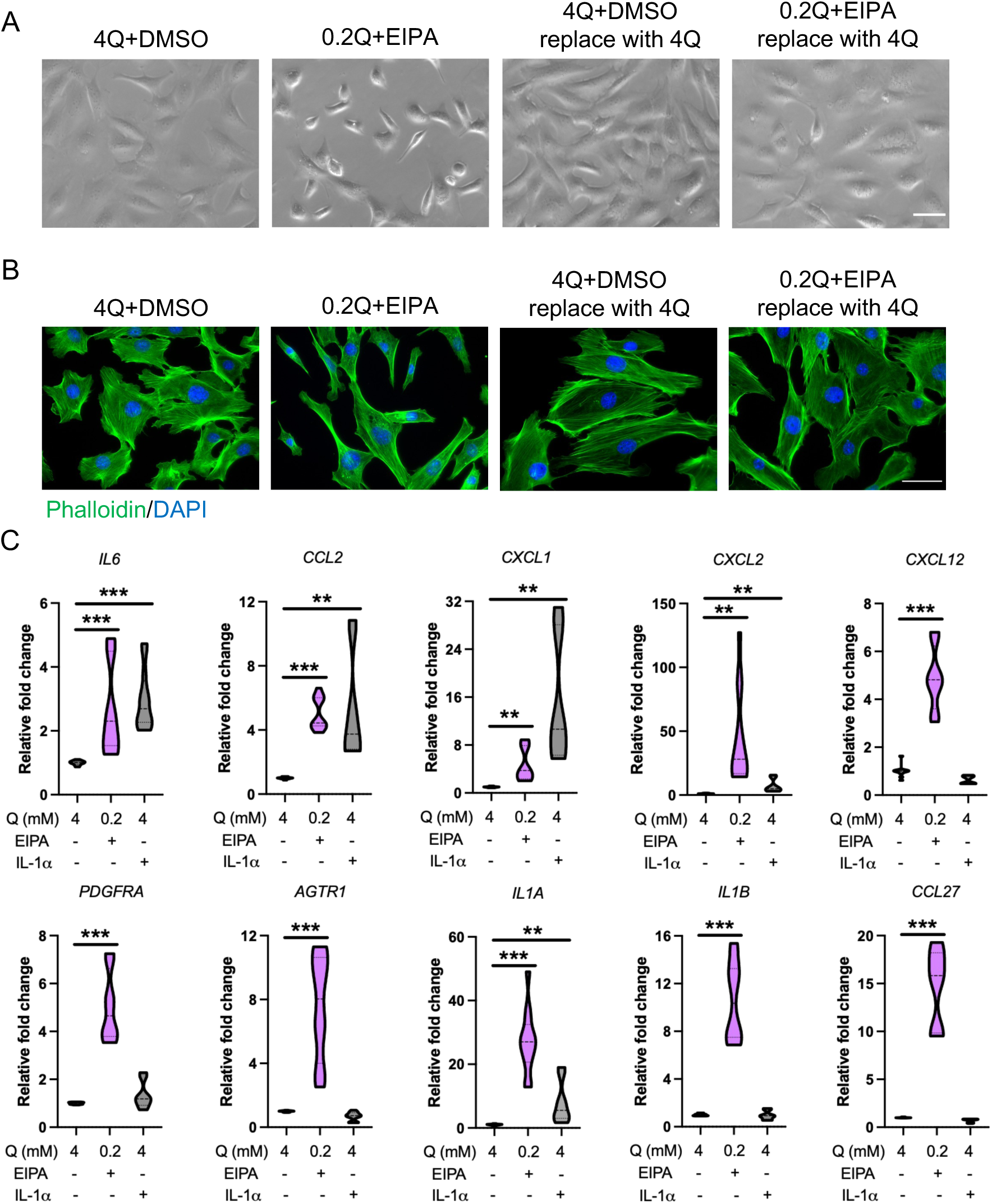
Characterization of the metabolic stress-induced intrinsic iCAF phenotype. (A) Mouse CAFs were treated with 4 mM Q with vehicle or 0.2 mM Q with EIPA. 32 hours post treatment, one set of samples were collected for RNA analysis as baseline and another set were washed with PBS and cultured with 4 mM Q medium for an additional 24 hours. Phase-contrast living cell images were captured prior to collection. Representative images are shown. Scale bar: 50 μm. (B) Cells were seeded on coverslips in 24-well format and treated as described above. Both sets of cells were washed with DPBS after treatment and fixed with 3.7% formaldehyde, followed by phalloidin (green) and DAPI (blue) staining. Representative images are shown. Scale bar: 50 μm. (C) Expression levels of the iCAF markers in CAFs treated with 0.2Q with 25 μM EIPA or 25 pg/ml murine IL-1α for 32 hours. 4mM Q was used as a reference control. Data from at least three independent experiments are shown as mean ± SEM, unpaired t-test. *p<0.05, **p<0.01, ***p<0.001.

**Supplemental Figure 4.**
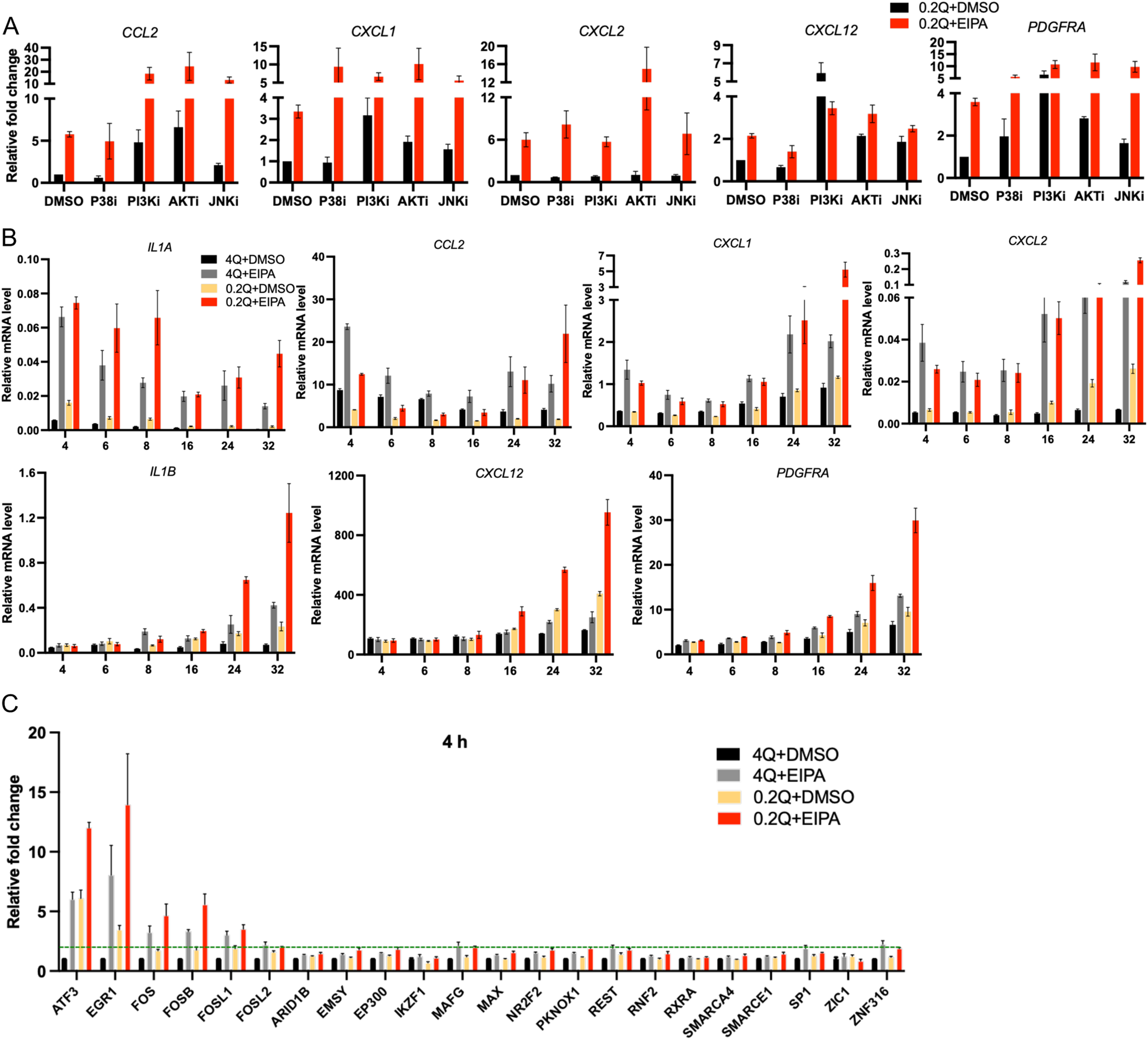
Regulation and dynamics of the metabolic stress-induced intrinsic iCAF phenotype. (A) Mouse CAFs were pre-treated with p38 inhibitor (P38i), PI3K inhibitor (PI3Ki), AKT inhibitor (AKTi), or JNK inhibitor (JNKi) for 1 hour and then treated with 4 mM or 0.2 mM Q with or without the presence of 25 μM EIPA for 24 hours. The mRNA levels of iCAF-related markers were analyzed by qPCR. (B) Mouse CAFs were treated with 4 mM or 0.2 mM Q with or without the presence of 25 μM EIPA for 4, 6, 8, 16, 24, and 32 hours. The mRNA levels of iCAF-related markers were analyzed by qPCR and normalized to RPL13. (C) Mouse CAFs were treated with 4 mM or 0.2 mM Q with or without the presence of 25 μM EIPA for 4 hours. The mRNA levels of 22 TFs were analyzed by qPCR.

**Supplemental Figure 5.**
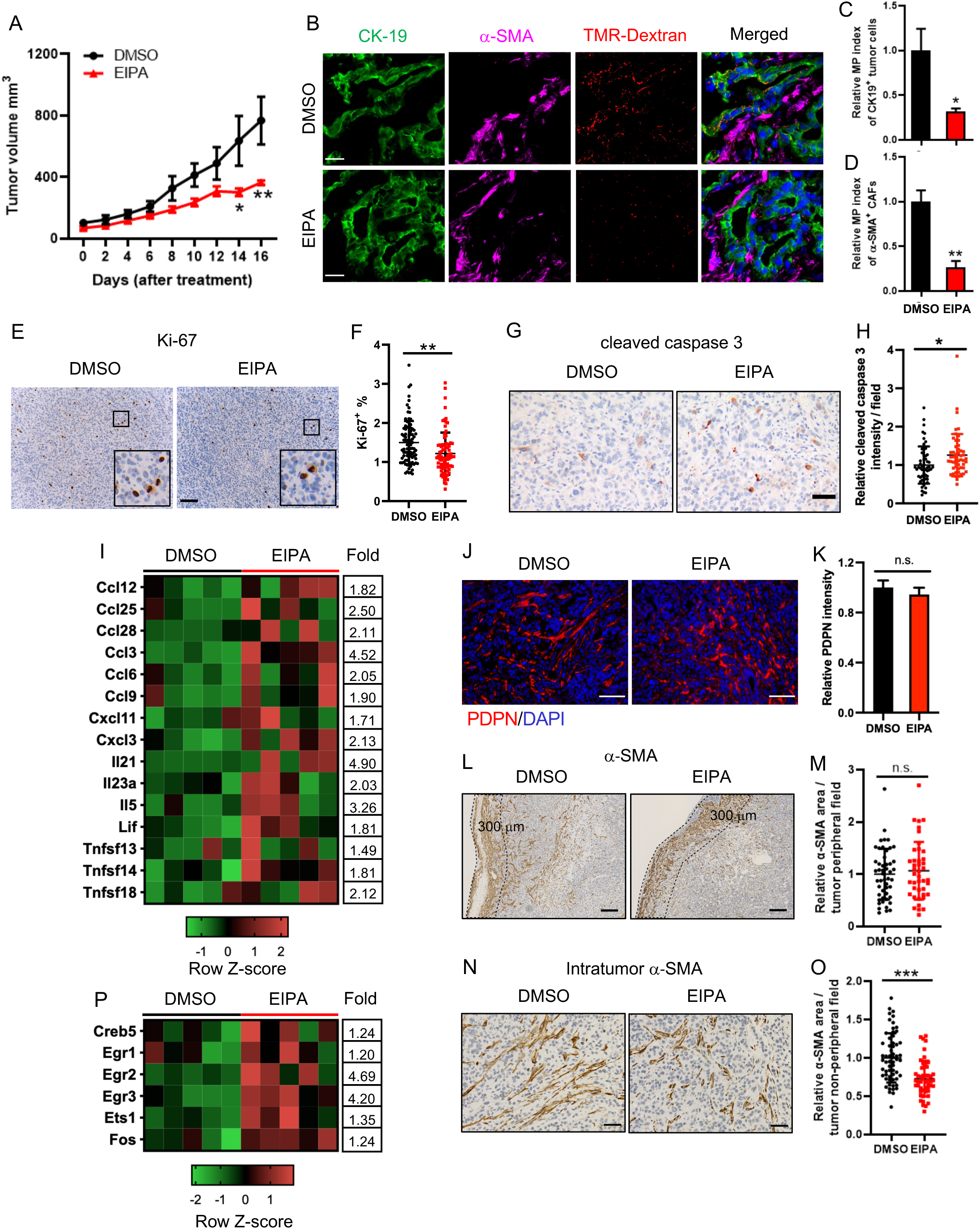
EIPA suppresses the growth of syngeneic heterotopic tumors and modulates proliferation and apoptosis in syngeneic orthotopic tumors. (A) Tumor growth curve of KPC heterotopic subcutaneous tumors treated with DMSO or EIPA at 10 mg/kg for two weeks. Data are shown as mean ± SEM (n=3 or 4). Two-way ANOVA. *p<0.05, **p<0.01. (B-D) Ex-vivo macropinocytic uptake of TMR-labeled dextran was performed on KPC heterotopic tumors treated with DMSO or EIPA. Tumor cells and myCAFs were labeled with CK-19 (green) or α-SMA (magenta), respectively (B). The relative macropinocytic index in the tumor cells (C) or the myCAFs (D) was computed. Data are shown as mean ± SEM (n=3), unpaired t-test. *p<0.05, **p<0.01. Scale bar: 20 μm. (E and F) Representative images (E) and quantification (F) of anti-Ki-67 staining in KPC orthotopic tumors treated with DMSO or EIPA. 79 or 88 fields (20x) for each group were analyzed. Data are shown as mean ± SD, unpaired t-test. **p<0.01. Scale bar: 100 μm. (G and H) Representative images (G) and quantification (H) of anti-cleaved caspase-3 staining in KPC orthotopic tumors treated with DMSO or EIPA. 52 or 56 fields (20x) for each group were analyzed. Data are shown as mean ± SD, unpaired t-test. *p<0.05. Scale bar: 50 μm. (I) Heatmap depicting gene expression of other inflammatory factors in DMSO (n=5) or EIPA (n=5)-treated PDAC tumors using nanoString nCounter mouse PanCancer Immune Profiling Panel. mRNA fold change for each gene is indicated. (J and K) Representative images (J) and quantification (K) of anti-podoplanin (PDPN) immunofluorescence staining in KPC orthotopic tumors treated with DMSO or EIPA. ∼120 fields from 6 tumors for each group were analyzed. Data are shown as mean ± SEM, unpaired t-test. n.s., not significant. Scale bar: 100 μm. (L-O) Anti-α-SMA staining. Representative images (L) and quantification (M) of α-SMA area at the periphery of KPC orthotopic tumors (<300μm from the edge of the tumor, outlined with black dashed line). 42 or 49 fields (5x) for each group were analyzed. Data are shown as mean ± SD, unpaired t-test. n.s., not significant. Scale bar: 200 μm. Representative images (N) and quantification (O) of α-SMA area at the intratumoral region of KPC orthotopic tumors. 60 fields (20x) from 6 DMSO-treated tumors and 50 fields (20x) from 5 EIPA-treated tumors were analyzed. Data are shown as mean ± SD, unpaired t-test. ***p<0.001. Scale bar: 50 μm. (P) Heatmap depicting gene expression of p-ERK1/2 targets in DMSO (n=5) or EIPA (n=5)-treated PDAC tumors using nanoString nCounter mouse PanCancer Immune Profiling Panel. mRNA fold change for each gene is indicated.

**Supplemental Figure 6.**
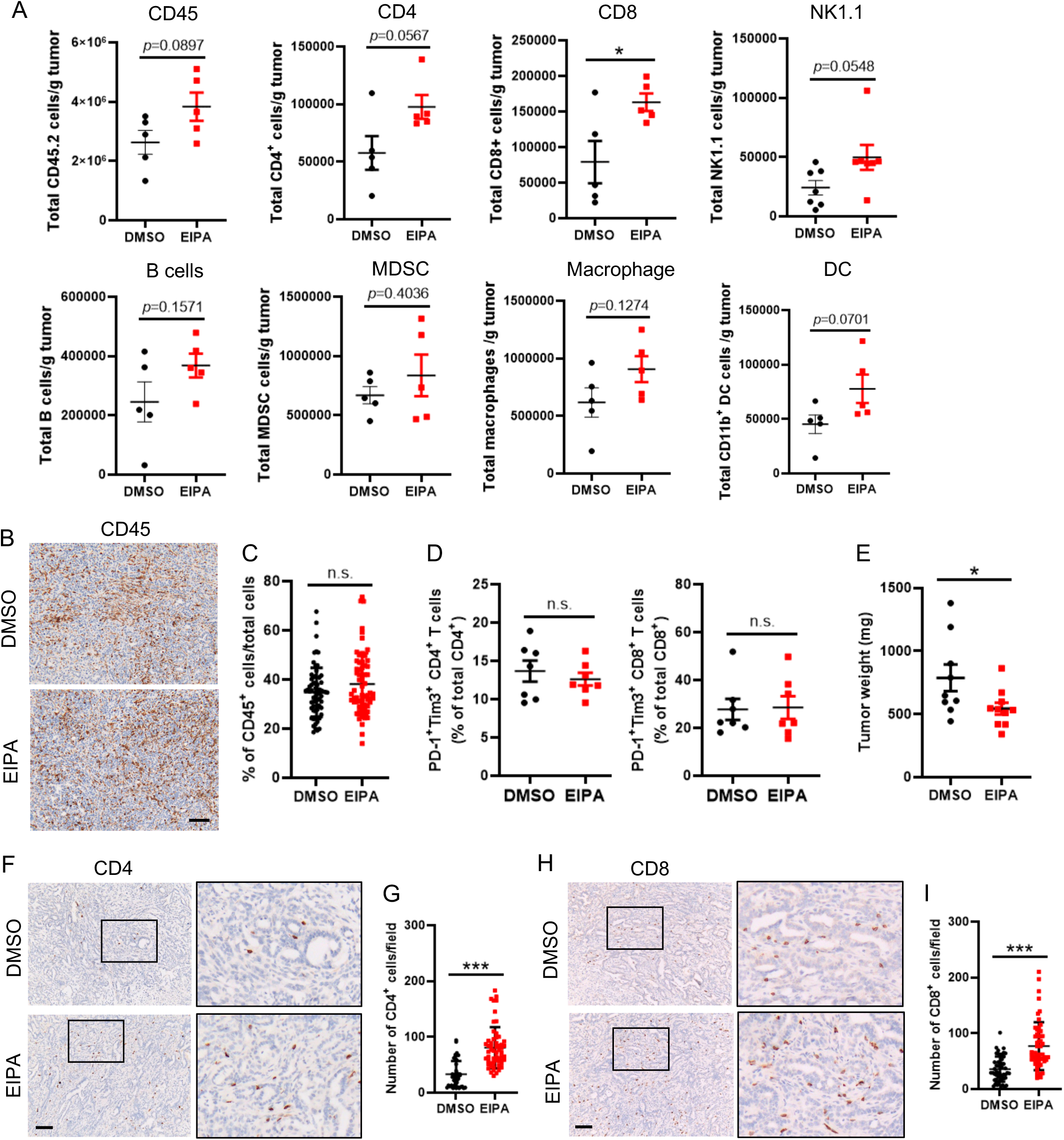
EIPA promotes T cell infiltration but its anti-tumor effects are not mediated by T cells. (A) Immune profiling was performed on dissociated KPC orthotopic tumors treated with DMSO or EIPA. The total number of each immune cell population was quantified using flow cytometry and expressed per gram of tumor. Data are shown as mean ± SEM, n=5 or 7, unpaired t-test. p values are as indicated. (B and C) Representative images (B) and quantification (C) of anti-CD45 staining in KPC orthotopic tumors treated with DMSO or EIPA. 64 or 66 fields (20x) for each group were analyzed. Data are shown as mean ± SD, unpaired t-test. n.s., not significant. Scale bar: 100 μm. (D) The percentage of CD4^+^ T cells or CD8^+^ T cells co-expressing PD-1 and Tim-3 in KPC orthotopic tumors treated with DMSO or EIPA. Data are shown as mean ± SEM, n=7, unpaired t-test. n.s., not significant. (E) KPC orthotopic tumors were generated in athymic nu/nu mice and treated with DMSO (n=9) or EIPA (n=10) for two weeks. Tumors were harvested and tumor weights were determined. Data are shown as mean ± SEM, unpaired t-test. *p<0.05. (F and G) Representative images (F) and quantification (G) of anti-CD4 staining in KPC heterotopic tumors treated with DMSO or EIPA. 43 or 56 fields (20x) for each group were analyzed. Data are shown as mean ± SD, unpaired t-test. **p<0.01. Scale bar: 100 μm. (H and I) Representative images (H) and quantification (I) of anti-CD8a staining in KPC heterotopic tumors treated with DMSO or EIPA. 49 or 55 fields (20x) for each group were analyzed. Data are shown as mean ± SD, unpaired t-test. *p<0.05. Scale bar: 100 μm.

**Supplemental Figure 7.**
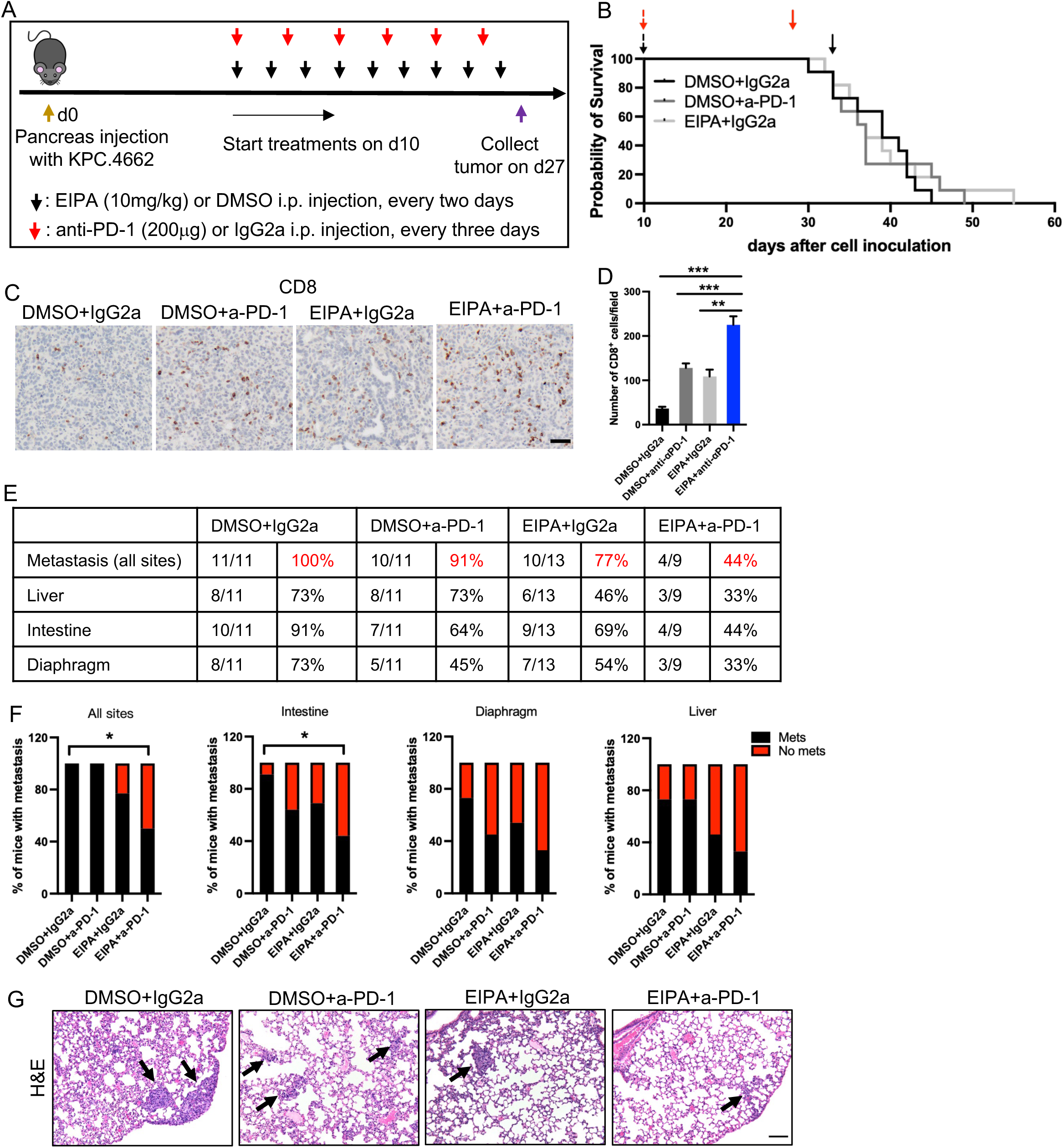
Combination treatment with EIPA and immunotherapy suppresses intestinal and lung metastasis. (A) Scheme depicting the treatment protocol for EIPA and anti-a-PD1 combination therapy. (B) Mice bearing orthotopic PDAC tumors were treated with EIPA or anti-PD-1 antibody alone with vehicle or isotype antibody as the control. Overall survival was analyzed using Kaplan-Meier curves with the log-rank test. (C and D) Representative images (C) and quantification (D) of anti-CD8a staining in KPC orthotopic tumors after treatment. Images were analyzed: DMSO+IgG2a, 54 fields (20x) from 4 tumors; EIPA+IgG2a, 41 fields (20x) from 5 tumors; DMSO+a-PD-1, 49 fields (20x) from 5 tumors; EIPA+a-PD-1, 43 fields (20x) from 4 tumors. Data are shown as mean ± SD, One-way ANOVA. ***p<0.001. Scale bar: 50 μm. (E and F) Tumor macrometastases to the liver, intestine and diaphragm were analyzed. The number of mice with macrometastases in each organ site was calculated and shown in the table (E). The percentage of mice with evidence of macrometastases to the intestine, liver, or diaphragm was determined (F). Chi-squared tests. *p<0.05. (G) Representative images of Haemotoxylin and Eosin (H&E) staining of mouse lungs after treatment. Arrows indicate micrometastases. Scale bar: 100 μm.

**Supplemental Figure 8.**
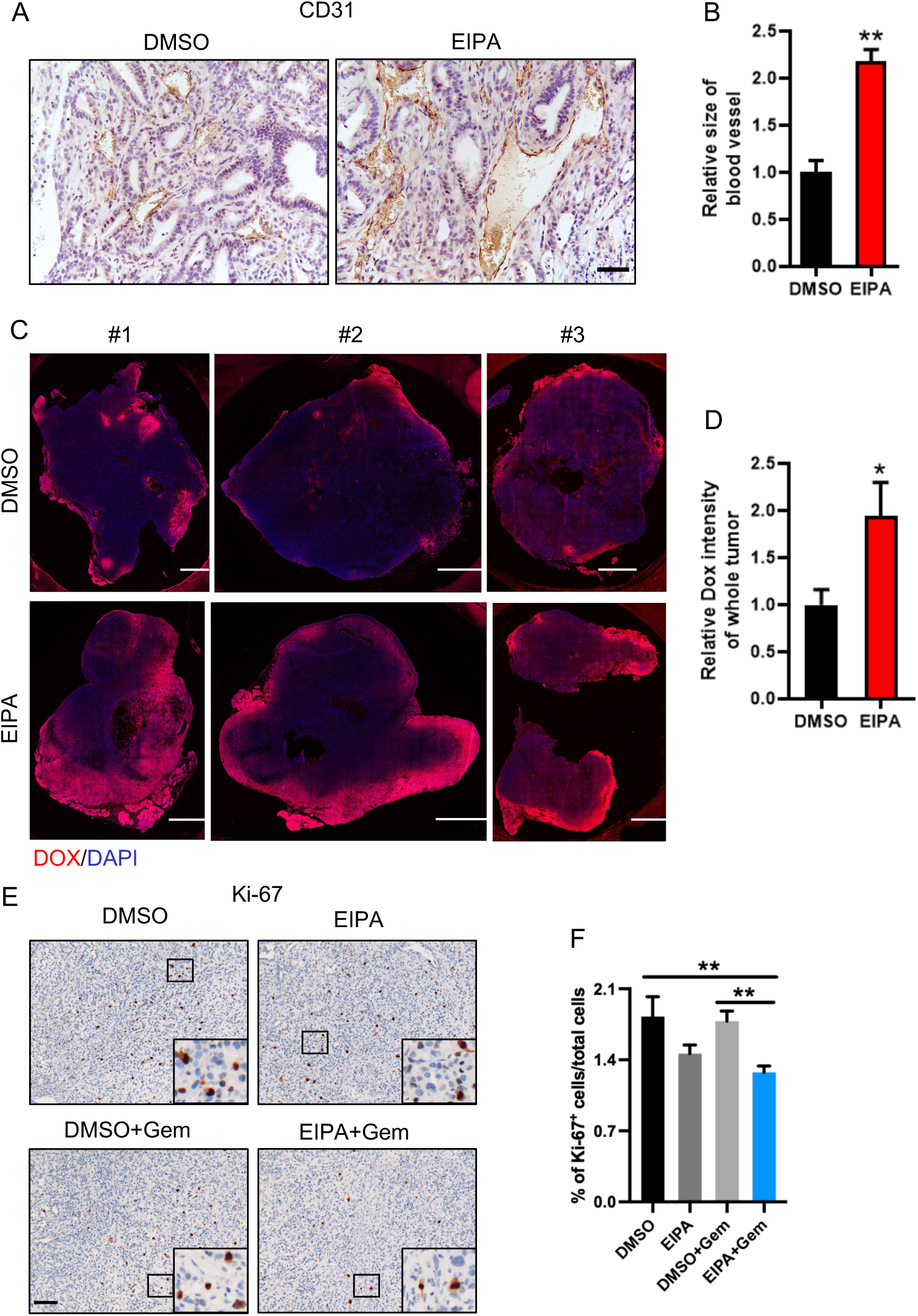
EIPA expands tumor vasculature, enhances drug delivery, and improves PDAC therapeutic responses in combination with gemcitabine. (A and B) Anti-CD31 staining in KPC heterotopic tumors treated with DMSO or EIPA. Representative images are shown (A). The area of CD31^+^ blood vessels in the tumor sections was computed for each field and is shown relative to the control (B). n=3 or 4. Scale bar: 50 μm. (C and D) Confocal microscope images of entire KPC orthotopic tumors that were treated with DOX after treatment with DMSO or EIPA (C). Mean fluorescent intensity of the DOX autofluorescence in the whole tumors was quantified and is shown relative to the control (D). Data are shown as mean ± SEM (n=9). Unpaired t-test. *p<0.05. (E and F) Representative images (E) and quantification (F) of anti-Ki-67 staining in KPC orthotopic tumors after treatment. Images were analyzed: DMSO, 53 fields (20x) from 5 tumors; EIPA, 70 fields (20x) from 6 tumors; DMSO+Gem, 85 fields (20x) from 5 tumors; EIPA+Gem, 88 fields (20x) from 6 tumors. Data are shown as mean ± SD, One-way ANOVA. **p<0.01, n.s., not significant. Scale bar: 100 μm.

## References

1. Sahai, E., Astsaturov, I., Cukierman, E., DeNardo, D.G., Egeblad, M., Evans, R.M., Fearon, D., Greten, F.R., Hingorani, S.R., Hunter, T., et al. (2020). A framework for advancing our understanding of cancer-associated fibroblasts. Nature reviews. Cancer 20, 174–186. 10.1038/s41568-019-0238-1.

2. LeBleu, V.S., and Kalluri, R. (2018). A peek into cancer-associated fibroblasts: origins, functions and translational impact. Dis Model Mech 11. ARTN dmm02944710.1242/dmm.029447.

3. Caligiuri, G., and Tuveson, D.A. (2023). Activated fibroblasts in cancer: Perspectives and challenges. Cancer cell 41, 434–449. 10.1016/j.ccell.2023.02.015.

4. Biffi, G., and Tuveson, D.A. (2021). Diversity and Biology of Cancer-Associated Fibroblasts. Physiological reviews 101, 147–176. 10.1152/physrev.00048.2019.

5. Sherman, M.H., and Beatty, G.L. (2023). Tumor Microenvironment in Pancreatic Cancer Pathogenesis and Therapeutic Resistance. Annual review of pathology 18, 123–148. 10.1146/annurev-pathmechdis-031621-024600.

6. Sousa, C.M., Biancur, D.E., Wang, X., Halbrook, C.J., Sherman, M.H., Zhang, L., Kremer, D., Hwang, R.F., Witkiewicz, A.K., Ying, H., et al. (2016). Pancreatic stellate cells support tumour metabolism through autophagic alanine secretion. Nature 536, 479–483. 10.1038/nature19084.

7. Zhang, Y., Recouvreux, M.V., Jung, M., Galenkamp, K.M.O., Li, Y., Zagnitko, O., Scott, D.A., Lowy, A.M., and Commisso, C. (2021). Macropinocytosis in Cancer-Associated Fibroblasts Is Dependent on CaMKK2/ARHGEF2 Signaling and Functions to Support Tumor and Stromal Cell Fitness. Cancer discovery 11, 1808–1825. 10.1158/2159-8290.CD-20-0119.

8. Francescone, R., Vendramini-Costa, D.B., Franco-Barraza, J., Wagner, J., Muir, A., Lau, A.N., Gabitova, L., Pazina, T., Gupta, S., Luong, T., et al. (2021). Netrin G1 Promotes Pancreatic Tumorigenesis through Cancer-Associated Fibroblast-Driven Nutritional Support and Immunosuppression. Cancer discovery 11, 446–479. 10.1158/2159-8290.CD-20-0775.

9. Liu, T., Han, C., Wang, S., Fang, P., Ma, Z., Xu, L., and Yin, R. (2019). Cancer-associated fibroblasts: an emerging target of anti-cancer immunotherapy. Journal of hematology & oncology 12, 86. 10.1186/s13045-019-0770-1.

10. Helms, E., Onate, M.K., and Sherman, M.H. (2020). Fibroblast Heterogeneity in the Pancreatic Tumor Microenvironment. Cancer discovery 10, 648–656. 10.1158/2159-8290.CD-19-1353.

11. Sherman, M.H., and Magliano, M.P.d. (2023). Cancer-Associated Fibroblasts: Lessons from Pancreatic Cancer. Annual Review of Cancer Biology 7, null. 10.1146/annurev-cancerbio-061421-035400.

12. McAndrews, K.M., Chen, Y., Darpolor, J.K., Zheng, X., Yang, S., Carstens, J.L., Li, B., Wang, H., Miyake, T., Correa de Sampaio, P., et al. (2022). Identification of Functional Heterogeneity of Carcinoma-Associated Fibroblasts with Distinct IL6-Mediated Therapy Resistance in Pancreatic Cancer. Cancer discovery 12, 1580–1597. 10.1158/2159-8290.CD-20-1484.

13. Vickman, R.E., Faget, D.V., Beachy, P., Beebe, D., Bhowmick, N.A., Cukierman, E., Deng, W.M., Granneman, J.G., Hildesheim, J., Kalluri, R., et al. (2020). Deconstructing tumor heterogeneity: the stromal perspective. Oncotarget 11, 3621–3632. 10.18632/oncotarget.27736.

14. Ohlund, D., Handly-Santana, A., Biffi, G., Elyada, E., Almeida, A.S., Ponz-Sarvise, M., Corbo, V., Oni, T.E., Hearn, S.A., Lee, E.J., et al. (2017). Distinct populations of inflammatory fibroblasts and myofibroblasts in pancreatic cancer. The Journal of experimental medicine 214, 579–596. 10.1084/jem.20162024.

15. Elyada, E., Bolisetty, M., Laise, P., Flynn, W.F., Courtois, E.T., Burkhart, R.A., Teinor, J.A., Belleau, P., Biffi, G., Lucito, M.S., et al. (2019). Cross-Species Single-Cell Analysis of Pancreatic Ductal Adenocarcinoma Reveals Antigen-Presenting Cancer-Associated Fibroblasts. Cancer discovery 9, 1102–1123. 10.1158/2159-8290.CD-19-0094.

16. Hutton, C., Heider, F., Blanco-Gomez, A., Banyard, A., Kononov, A., Zhang, X.H., Karim, S., Paulus-Hock, V., Watt, D., Steele, N., et al. (2021). Single-cell analysis defines a pancreatic fibroblast lineage that supports anti-tumor immunity. Cancer cell 39, 1227-+. 10.1016/j.ccell.2021.06.017.

17. Helms, E.J., Berry, M.W., Chaw, R.C., DuFort, C.C., Sun, D., Onate, M.K., Oon, C., Bhattacharyya, S., Sanford-Crane, H., Horton, W., et al. (2022). Mesenchymal Lineage Heterogeneity Underlies Nonredundant Functions of Pancreatic Cancer-Associated Fibroblasts. Cancer discovery 12, 484–501. 10.1158/2159-8290.CD-21-0601.

18. Huang, H., Wang, Z., Zhang, Y., Pradhan, R.N., Ganguly, D., Chandra, R., Murimwa, G., Wright, S., Gu, X., Maddipati, R., et al. (2022). Mesothelial cell-derived antigen-presenting cancer-associated fibroblasts induce expansion of regulatory T cells in pancreatic cancer. Cancer cell 40, 656–673 e657. 10.1016/j.ccell.2022.04.011.

19. Ozdemir, B.C., Pentcheva-Hoang, T., Carstens, J.L., Zheng, X., Wu, C.C., Simpson, T.R., Laklai, H., Sugimoto, H., Kahlert, C., Novitskiy, S.V., et al. (2014). Depletion of carcinoma-associated fibroblasts and fibrosis induces immunosuppression and accelerates pancreas cancer with reduced survival. Cancer cell 25, 719–734. 10.1016/j.ccr.2014.04.005.

20. Biffi, G., Oni, T.E., Spielman, B., Hao, Y., Elyada, E., Park, Y., Preall, J., and Tuveson, D.A. (2019). IL1-Induced JAK/STAT Signaling Is Antagonized by TGF beta to Shape CAF Heterogeneity in Pancreatic Ductal Adenocarcinoma. Cancer discovery 9, 282–301. 10.1158/2159-8290.CD-18-0710.

21. Steele, N.G., Biffi, G., Kemp, S.B., Zhang, Y., Drouillard, D., Syu, L., Hao, Y., Oni, T.E., Brosnan, E., Elyada, E., et al. (2021). Inhibition of Hedgehog Signaling Alters Fibroblast Composition in Pancreatic Cancer. Clinical cancer research : an official journal of the American Association for Cancer Research 27, 2023–2037. 10.1158/1078-0432.CCR-20-3715.

22. Mello, A.M., Ngodup, T., Lee, Y., Donahue, K.L., Li, J., Rao, A., Carpenter, E.S., Crawford, H.C., Pasca di Magliano, M., and Lee, K.E. (2022). Hypoxia promotes an inflammatory phenotype of fibroblasts in pancreatic cancer. Oncogenesis 11, 56. 10.1038/s41389-022-00434-2.

23. Schwoerer, S., Cimino, F.V., Ros, M., Tsanov, K.M., Ng, C., Lowe, S.W., Carmona-Fontaine, C., and Thompson, C.B. (2023). Hypoxia potentiates the inflammatory fibroblast phenotype promoted by pancreatic cancer cell-derived cytokines. Cancer research. 10.1158/0008-5472.CAN-22-2316.

24. Singh, S.P., Dosch, A.R., Mehra, S., De Castro Silva, I., Bianchi, A., Garrido, V.T., Zhou, Z., Adams, A., Amirian, H., Box, E.W., et al. (2024). Tumor cell-intrinsic p38 MAPK signaling promotes IL1⍺-mediated stromal inflammation and therapeutic resistance in pancreatic cancer. Cancer research. 10.1158/0008-5472.CAN-23-1200.

25. Recouvreux, M.V., Moldenhauer, M.R., Galenkamp, K.M.O., Jung, M., James, B., Zhang, Y., Lowy, A., Bagchi, A., and Commisso, C. (2020). Glutamine depletion regulates Slug to promote EMT and metastasis in pancreatic cancer. The Journal of experimental medicine 217. 10.1084/jem.20200388.

26. Lee, S.W., Zhang, Y., Jung, M., Cruz, N., Alas, B., and Commisso, C. (2019). EGFR-Pak Signaling Selectively Regulates Glutamine Deprivation-Induced Macropinocytosis. Developmental cell 50, 381–392 e385. 10.1016/j.devcel.2019.05.043.

27. Kamphorst, J.J., Nofal, M., Commisso, C., Hackett, S.R., Lu, W., Grabocka, E., Vander Heiden, M.G., Miller, G., Drebin, J.A., Bar-Sagi, D., et al. (2015). Human pancreatic cancer tumors are nutrient poor and tumor cells actively scavenge extracellular protein. Cancer research 75, 544–553. 75/3/544 [pii]10.1158/0008-5472.CAN-14-2211.

28. Commisso, C., Davidson, S.M., Soydaner-Azeloglu, R.G., Parker, S.J., Kamphorst, J.J., Hackett, S., Grabocka, E., Nofal, M., Drebin, J.A., Thompson, C.B., et al. (2013). Macropinocytosis of protein is an amino acid supply route in Ras-transformed cells. Nature 497, 633–637. nature12138 [pii]10.1038/nature12138.

29. Davidson, S.M., Jonas, O., Keibler, M.A., Hou, H.W., Luengo, A., Mayers, J.R., Wyckoff, J., Del Rosario, A.M., Whitman, M., Chin, C.R., et al. (2017). Direct evidence for cancer-cell-autonomous extracellular protein catabolism in pancreatic tumors. Nature medicine 23, 235–241. 10.1038/nm.4256.

30. Sullivan, M.R., Danai, L.V., Lewis, C.A., Chan, S.H., Gui, D.Y., Kunchok, T., Dennstedt, E.A., Vander Heiden, M.G., and Muir, A. (2019). Quantification of microenvironmental metabolites in murine cancers reveals determinants of tumor nutrient availability. Elife 8. 10.7554/eLife.44235.

31. Ye, J.B., Palm, W., Peng, M., King, B., Lindsten, T., Li, M.O., Koumenis, C., and Thompson, C.B. (2015). GCN2 sustains mTORC1 suppression upon amino acid deprivation by inducing Sestrin2. Genes & development 29, 2331–2336. 10.1101/gad.269324.115.

32. Kennel, K.B., Bozlar, M., De Valk, A.F., and Greten, F.R. (2023). Cancer-Associated Fibroblasts in Inflammation and Antitumor Immunity. Clinical cancer research : an official journal of the American Association for Cancer Research 29, 1009–1016. 10.1158/1078-0432.CCR-22-1031.

33. Schworer, S., Cimino, F.V., Ros, M., Tsanov, K.M., Ng, C., Lowe, S.W., Carmona-Fontaine, C., and Thompson, C.B. (2023). Hypoxia Potentiates the Inflammatory Fibroblast Phenotype Promoted by Pancreatic Cancer Cell-Derived Cytokines. Cancer research 83, 1596–1610. 10.1158/0008-5472.CAN-22-2316.

34. Encarnacion-Rosado, J., Sohn, A.S.W., Biancur, D.E., Lin, E.Y., Osorio-Vasquez, V., Rodrick, T., Gonzalez-Baerga, D., Zhao, E., Yokoyama, Y., Simeone, D.M., et al. (2024). Targeting pancreatic cancer metabolic dependencies through glutamine antagonism. Nat Cancer 5, 85–99. 10.1038/s43018-023-00647-3.

35. Fishilevich, S., Nudel, R., Rappaport, N., Hadar, R., Plaschkes, I., Iny Stein, T., Rosen, N., Kohn, A., Twik, M., Safran, M., et al. (2017). GeneHancer: genome-wide integration of enhancers and target genes in GeneCards. Database (Oxford) 2017. 10.1093/database/bax028.

36. Su, H., Yang, F., Fu, R., Li, X., French, R., Mose, E., Pu, X., Trinh, B., Kumar, A., Liu, J., et al. (2021). Cancer cells escape autophagy inhibition via NRF2-induced macropinocytosis. Cancer cell 39, 678–693 e611. 10.1016/j.ccell.2021.02.016.

37. Kim, S.M., Nguyen, T.T., Ravi, A., Kubiniok, P., Finicle, B.T., Jayashankar, V., Malacrida, L., Hou, J., Robertson, J., Gao, D., et al. (2018). PTEN Deficiency and AMPK Activation Promote Nutrient Scavenging and Anabolism in Prostate Cancer Cells. Cancer discovery 8, 866–883. 10.1158/2159-8290.CD-17-1215.

38. Bayne, L.J., Beatty, G.L., Jhala, N., Clark, C.E., Rhim, A.D., Stanger, B.Z., and Vonderheide, R.H. (2012). Tumor-derived granulocyte-macrophage colony-stimulating factor regulates myeloid inflammation and T cell immunity in pancreatic cancer. Cancer cell 21, 822–835. 10.1016/j.ccr.2012.04.025.

39. Vennin, C., Murphy, K.J., Morton, J.P., Cox, T.R., Pajic, M., and Timpson, P. (2018). Reshaping the Tumor Stroma for Treatment of Pancreatic Cancer. Gastroenterology 154, 820–838. 10.1053/j.gastro.2017.11.280.

40. Ene-Obong, A., Clear, A.J., Watt, J., Wang, J., Fatah, R., Riches, J.C., Marshall, J.F., Chin-Aleong, J., Chelala, C., Gribben, J.G., et al. (2013). Activated pancreatic stellate cells sequester CD8+ T cells to reduce their infiltration of the juxtatumoral compartment of pancreatic ductal adenocarcinoma. Gastroenterology 145, 1121–1132. 10.1053/j.gastro.2013.07.025.

41. Froeling, F.E., Feig, C., Chelala, C., Dobson, R., Mein, C.E., Tuveson, D.A., Clevers, H., Hart, I.R., and Kocher, H.M. (2011). Retinoic acid-induced pancreatic stellate cell quiescence reduces paracrine Wnt-beta-catenin signaling to slow tumor progression. Gastroenterology 141, 1486–1497, 1497 e1481-1414. 10.1053/j.gastro.2011.06.047.

42. Saka, D., Gokalp, M., Piyade, B., Cevik, N.C., Arik Sever, E., Unutmaz, D., Ceyhan, G.O., Demir, I.E., and Asimgil, H. (2020). Mechanisms of T-Cell Exhaustion in Pancreatic Cancer. Cancers 12. 10.3390/cancers12082274.

43. Li, J., Byrne, K.T., Yan, F., Yamazoe, T., Chen, Z., Baslan, T., Richman, L.P., Lin, J.H., Sun, Y.H., Rech, A.J., et al. (2018). Tumor Cell-Intrinsic Factors Underlie Heterogeneity of Immune Cell Infiltration and Response to Immunotherapy. Immunity 49, 178–193 e177. 10.1016/j.immuni.2018.06.006.

44. Provenzano, P.P., Cuevas, C., Chang, A.E., Goel, V.K., Von Hoff, D.D., and Hingorani, S.R. (2012). Enzymatic targeting of the stroma ablates physical barriers to treatment of pancreatic ductal adenocarcinoma. Cancer cell 21, 418–429. 10.1016/j.ccr.2012.01.007.

45. Jacobetz, M.A., Chan, D.S., Neesse, A., Bapiro, T.E., Cook, N., Frese, K.K., Feig, C., Nakagawa, T., Caldwell, M.E., Zecchini, H.I., et al. (2013). Hyaluronan impairs vascular function and drug delivery in a mouse model of pancreatic cancer. Gut 62, 112–120. 10.1136/gutjnl-2012-302529.

46. Sherman, M.H., Yu, R.T., Engle, D.D., Ding, N., Atkins, A.R., Tiriac, H., Collisson, E.A., Connor, F., Van Dyke, T., Kozlov, S., et al. (2014). Vitamin D receptor-mediated stromal reprogramming suppresses pancreatitis and enhances pancreatic cancer therapy. Cell 159, 80–93. 10.1016/j.cell.2014.08.007.

47. Duarte, D., and Vale, N. (2022). Evaluation of synergism in drug combinations and reference models for future orientations in oncology. Curr Res Pharmacol Drug Discov 3, 100110. 10.1016/j.crphar.2022.100110.

48. Jayashankar, V., and Edinger, A.L. (2020). Macropinocytosis confers resistance to therapies targeting cancer anabolism. Nature communications 11, 1121. 10.1038/s41467-020-14928-3.

49. Mishra, R., Haldar, S., Placencio, V., Madhav, A., Rohena-Rivera, K., Agarwal, P., Duong, F., Angara, B., Tripathi, M., Liu, Z., et al. (2018). Stromal epigenetic alterations drive metabolic and neuroendocrine prostate cancer reprogramming. J Clin Invest 128, 4472–4484. 10.1172/JCI99397.

50. Romer, A.M.A., Thorseth, M.L., and Madsen, D.H. (2021). Immune Modulatory Properties of Collagen in Cancer. Frontiers in immunology 12, 791453. 10.3389/fimmu.2021.791453.

51. Kuczek, D.E., Larsen, A.M.H., Thorseth, M.L., Carretta, M., Kalvisa, A., Siersbaek, M.S., Simoes, A.M.C., Roslind, A., Engelholm, L.H., Noessner, E., et al. (2019). Collagen density regulates the activity of tumor-infiltrating T cells. Journal for immunotherapy of cancer 7, 68. 10.1186/s40425-019-0556-6.

52. Sun, X., Wu, B., Chiang, H.C., Deng, H., Zhang, X., Xiong, W., Liu, J., Rozeboom, A.M., Harris, B.T., Blommaert, E., et al. (2021). Tumour DDR1 promotes collagen fibre alignment to instigate immune exclusion. Nature 599, 673–678. 10.1038/s41586-021-04057-2.

53. Nicolas-Boluda, A., Vaquero, J., Vimeux, L., Guilbert, T., Barrin, S., Kantari-Mimoun, C., Ponzo, M., Renault, G., Deptula, P., Pogoda, K., et al. (2021). Tumor stiffening reversion through collagen crosslinking inhibition improves T cell migration and anti-PD-1 treatment. Elife 10. ARTN e5868810.7554/eLife.58688.

54. Maddalena, M., Mallel, G., Nataraj, N.B., Shreberk-Shaked, M., Hassin, O., Mukherjee, S., Arandkar, S., Rotkopf, R., Kapsack, A., Lambiase, G., et al. (2021). TP53 missense mutations in PDAC are associated with enhanced fibrosis and an immunosuppressive microenvironment. Proceedings of the National Academy of Sciences of the United States of America 118. 10.1073/pnas.2025631118.

55. Rhim, A.D., Oberstein, P.E., Thomas, D.H., Mirek, E.T., Palermo, C.F., Sastra, S.A., Dekleva, E.N., Saunders, T., Becerra, C.P., Tattersall, I.W., et al. (2014). Stromal elements act to restrain, rather than support, pancreatic ductal adenocarcinoma. Cancer cell 25, 735–747. 10.1016/j.ccr.2014.04.021.

56. Kieffer, Y., Hocine, H.R., Gentric, G., Pelon, F., Bernard, C., Bourachot, B., Lameiras, S., Albergante, L., Bonneau, C., Guyard, A., et al. (2020). Single-Cell Analysis Reveals Fibroblast Clusters Linked to Immunotherapy Resistance in Cancer. Cancer discovery 10, 1330–1351. 10.1158/2159-8290.CD-19-1384.

57. Kay, E.J., and Zanivan, S. (2021). Metabolic pathways fuelling protumourigenic cancer-associated fibroblast functions. Curr Opin Syst Biol 28. 10.1016/j.coisb.2021.100377.

58. Eckert, M.A., Coscia, F., Chryplewicz, A., Chang, J.W., Hernandez, K.M., Pan, S., Tienda, S.M., Nahotko, D.A., Li, G., Blazenovic, I., et al. (2019). Proteomics reveals NNMT as a master metabolic regulator of cancer-associated fibroblasts. Nature 569, 723-+. 10.1038/s41586-019-1173-8.

59. Kay, E.J., Paterson, K., Riera-Domingo, C., Sumpton, D., Dabritz, J.H.M., Tardito, S., Boldrini, C., Hernandez-Fernaud, J.R., Athineos, D., Dhayade, S., et al. (2022). Cancer-associated fibroblasts require proline synthesis by PYCR1 for the deposition of pro-tumorigenic extracellular matrix. Nat Metab 4, 693–710. 10.1038/s42255-022-00582-0.

60. DuFort, C.C., DelGiorno, K.E., Carlson, M.A., Osgood, R.J., Zhao, C.M., Huang, Z.D., Thompson, C.B., Connor, R.J., Thanos, C.D., Brockenbrough, J.S., et al. (2016). Interstitial Pressure in Pancreatic Ductal Adenocarcinoma Is Dominated by a Gel-Fluid Phase. Biophys J 110, 2106–2119. 10.1016/j.bpj.2016.03.040.

61. Nia, H.T., Munn, L.L., and Jain, R.K. (2020). Physical traits of cancer. Science 370. 10.1126/science.aaz0868.

62. Olive, K.P., Jacobetz, M.A., Davidson, C.J., Gopinathan, A., McIntyre, D., Honess, D., Madhu, B., Goldgraben, M.A., Caldwell, M.E., Allard, D., et al. (2009). Inhibition of Hedgehog signaling enhances delivery of chemotherapy in a mouse model of pancreatic cancer. Science 324, 1457–1461. 10.1126/science.1171362.

63. Catenacci, D.V., Junttila, M.R., Karrison, T., Bahary, N., Horiba, M.N., Nattam, S.R., Marsh, R., Wallace, J., Kozloff, M., Rajdev, L., et al. (2015). Randomized Phase Ib/II Study of Gemcitabine Plus Placebo or Vismodegib, a Hedgehog Pathway Inhibitor, in Patients With Metastatic Pancreatic Cancer. Journal of clinical oncology : official journal of the American Society of Clinical Oncology 33, 4284–4292. 10.1200/JCO.2015.62.8719.

64. De Jesus-Acosta, A., Sugar, E.A., O’Dwyer, P.J., Ramanathan, R.K., Von Hoff, D.D., Rasheed, Z., Zheng, L., Begum, A., Anders, R., Maitra, A., et al. (2020). Phase 2 study of vismodegib, a hedgehog inhibitor, combined with gemcitabine and nab-paclitaxel in patients with untreated metastatic pancreatic adenocarcinoma. British journal of cancer 122, 498–505. 10.1038/s41416-019-0683-3.

65. Kim, E.J., Sahai, V., Abel, E.V., Griffith, K.A., Greenson, J.K., Takebe, N., Khan, G.N., Blau, J.L., Craig, R., Balis, U.G., et al. (2014). Pilot clinical trial of hedgehog pathway inhibitor GDC-0449 (vismodegib) in combination with gemcitabine in patients with metastatic pancreatic adenocarcinoma. Clinical cancer research : an official journal of the American Association for Cancer Research 20, 5937–5945. 10.1158/1078-0432.CCR-14-1269.

66. Tiwari, A., Tashiro, K., Dixit, A., Soni, A., Vogel, K., Hall, B., Shafqat, I., Slaughter, J., Param, N., Le, A., et al. (2020). Loss of HIF1A From Pancreatic Cancer Cells Increases Expression of PPP1R1B and Degradation of p53 to Promote Invasion and Metastasis. Gastroenterology 159, 1882–1897 e1885. 10.1053/j.gastro.2020.07.046.

67. Commisso, C., Flinn, R.J., and Bar-Sagi, D. (2014). Determining the macropinocytic index of cells through a quantitative image-based assay. Nature Protocols 9, 182–192. 10.1038/nprot.2014.004.

68. Galenkamp, K.M.O., Alas, B., and Commisso, C. (2019). Quantitation of Macropinocytosis in Cancer Cells. Methods in molecular biology 1928, 113–123. 10.1007/978-1-4939-9027-6_8.

69. Galenkamp, K.M.O., Galapate, C.M., Zhang, Y., and Commisso, C. (2021). Automated Imaging and Analysis for the Quantification of Fluorescently Labeled Macropinosomes. Journal of visualized experiments : JoVE. 10.3791/62828.

70. Lee, S.W., Alas, B., and Commisso, C. (2019). Detection and Quantification of Macropinosomes in Pancreatic Tumors. Methods in molecular biology 1882, 171–181. 10.1007/978-1-4939-8879-2_16.

